# Quantitative analysis and genome-scale modeling of human CD4^+^ T-cell differentiation reveals subset-specific regulation of glycosphingolipid pathways

**DOI:** 10.1101/2021.01.29.428853

**Authors:** Partho Sen, Syed Bilal Ahmad Andrabi, Tanja Buchacher, Mohd Moin Khan, Ubaid Ullah, Tuomas Lindeman, Marina Alves Amaral, Victoria Hinkkanen, Esko Kemppainen, Alex M Dickens, Omid Rasool, Tuulia Hyötyläinen, Riitta Lahesmaa, Matej Orešič

## Abstract

T-cells are sentinels of adaptive cell-mediated immune responses. T-cell activation, proliferation and differentiation involves metabolic reprogramming involving the interplay of genes, proteins and metabolites. Here, we aim to understand the metabolic pathways involved in the activation and functional differentiation of human CD4^+^ T-cell subsets (Th1, Th2, Th17 and iTregs). We combined genome-scale metabolic modeling, gene expression data, targeted and non-targeted lipidomics experiments, together with *in vitro* gene knockdown experiments and showed that human CD4^+^ T-cells undergo specific metabolic changes during activation and functional differentiation. In addition, we identified and confirmed the importance of ceramide and glycosphingolipid synthesis pathways in Th17 differentiation and effector functions. Finally, through *in vitro* gene knockdown experiments, we substantiated the requirement of serine palmitoyl transferase (*SPT*), a *de novo* sphingolipid pathway in the expression of proinflammatory cytokine (IL17A and IL17F) by Th17 cells. Our findings may provide a comprehensive resource for identifying CD4^+^ T-cell-specific targets for their selective manipulation under disease conditions, particularly, diseases characterized by an imbalance of Treg / Th17 cells. Our data also suggest a role for elevated levels of ceramides in conditions comorbid with these diseases, *e.g.*, obesity and insulin resistance.

## INTRODUCTION

CD4^+^ T-cells play a central role in the adaptive immune system. They orchestrate the immune responses and mediate protective immunity against pathogens [1]. An aberrant T-cell response is associated with cancer and autoimmune disorders [2, 3]. Circulating naïve T helper (Th) cells are metabolically quiescent and predominantly use oxidative phosphorylation (OXPHOS) to fuel their biological processes [4–7]. When exposed to antigens, naïve T-cells undergo activation, clonal expansion and differentiation to various effector (T_eff_) cells including Th1, Th2, Th17 and regulatory T-cells (Tregs), each driving various aspects of the immune responses [8, 9].

Upon activation, T-cells undergo metabolic reprogramming in order to provide energy and biosynthetic intermediates for growth and effector functions [5, 7, 10–12]. At this stage, aerobic glycolysis is augmented (the Warburg effect), which increases the activities of glycolytic enzymes. Their extracellular uptake of glucose increases by 40-50% [13], which, in turn, enhances their lactate production. Concomitantly, oxygen (O2) intake is increased by ~60% [14], whilst the utilization of glucose *via* OXPHOS is reduced [4]. Activated T-cells induces a metabolic sensor, *i.e.*, mammalian target of rapamycin (mTOR), to either differentiate into T_eff_ cells, or become suppressive Treg cells [15]. mTOR signaling induces the transcription factors Myc and HIF-1α, driving the expression of genes important for glycolysis and glutaminolysis as well as regulate STAT signaling for T-cell differentiation [16].

There is increasing evidence that metabolic reprograming may convert the proinflammatory Th17 phenotype towards the anti-inflammatory Treg phenotype in pathological disease condition [15, 17]. Furthermore, dysregulation of such metabolic reprogramming of T-cells can impair their clonal expansion [6, 7]. Depletion of glutamine in *in vitro* culture markedly impairs proliferation and cytokine production of T-cells [18]. Also, increased intracellular L-arginine is linked to metabolic regulation, survival and the anti-tumor activity of T-cells [19]. T-cell subsets such as Th1, Th2, and Th17, require acetyl-CoA carboxylase I (ACC1) for maturation [5]. In mice, T-cell-specific ACC1 deletion, or inhibition by inhibitor Soraphen A, prevents Th17 differentiation and cell-mediated autoimmune disease development [20].

Taken together, there is clear evidence that T-cell activation, differentiation and effector function is intrinsically linked to metabolic pathways [5, 7, 10–12, 21–23]. However, not much is known about the common and specific metabolic signatures, in human CD4^+^ T-cells and their functional subsets. Such knowledge could enable the selective manipulation of metabolism in these cells, with relevance to specific disease conditions [24].

Genome-scale metabolic models (GEMs) are computational frameworks which link the genes, proteins / enzymes, metabolites and pathways found in cells, tissues, organs, and organisms [25–28]. Over the past decade, genome-scale metabolic modeling (GSMM) has emerged as a powerful tool to study metabolism in human cells [26, 29, 30]. GSMM allows us to infer mechanistic relationship between genotype and phenotype [25–28].

Here we combined GSMM, published gene expression data, targeted and non-targeted lipidomics experiments, together with *in vitro* gene silencing, with the aim of understanding how human CD4^+^ T-cells modulate their metabolism during activation and subsequent functional differentiation. Our integrative approach identified several metabolic processes of interest, namely, we reveal the essentiality of glycosphingolipid (GSL) pathways in Th17 cells.

## RESULTS

### HTimmR: a genome-scale metabolic reconstruction of human CD4^+^ T-cells

We developed ‘Human T-immuno reconstructor’ (HTimmR), a generic and consensus metabolic reconstruction of human CD4^+^ T-cells. HTimmR includes 3841 metabolic genes (MGs), 7558 reactions, and 5140 metabolites (**Figure 1A**). HTimmR includes eight cellular compartments: extracellular cavity, peroxisome, mitochondria, cytosol, lysosome, endoplasmic reticulum, golgi apparatus, nucleus, and a cellular boundary which mimics a CD4^+^ T-cell. HTimmR was contextualized, *i.e.*, the active metabolic reactions in the model were selected using the gene expression data (**Methods**). Cell-type functional GEMs for T-naïve (Thp), Th1, Th2, Th17 and iTreg cells were developed (**Figure 1A**). The genes, reactions, and metabolites of these GEMs are given in (**Figure S1**).

**Figure 1.**
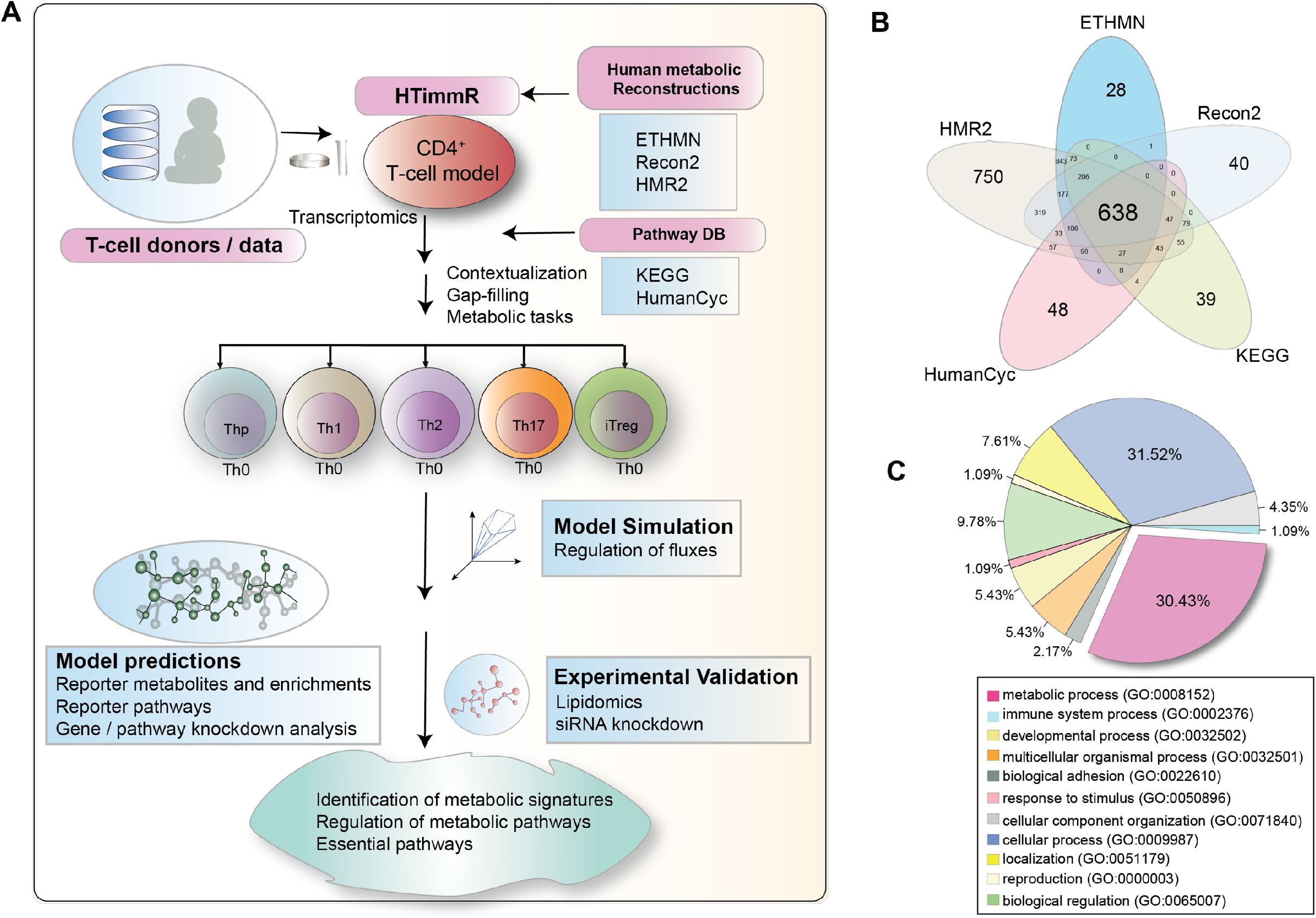
Metabolic reconstruction of human CD4^+^ T-cell subsets. **(A)** A schematic representation showing metabolic reconstruction of generic CD4^+^ T-cell (HTimmR), and contextualization of HTimmR to functional genome-scale models (GEMs) for T-naïve (Thp), T-activated (Th0), and differentiated T-helper (Th) subsets, using lineage-specific gene expression datasets. (**B**) Venn diagram showing metabolic genes (MGs) of human CD4^+^ T-cells identified in this study, which were commonly or uniquely found in various human metabolic reconstructions (HMR2, ETHMN, RECON), and pathway databases (KEGG and humanCyc). (**C**) A pie-chart showing the gene ontology (GO term) mapping of several biological processes exhibited by the MGs of the CD4^+^ T-cells.

### Identification of metabolic genes of human CD4^+^ T-cell activation and subsets differentiation

When mapping the published gene expression data of each CD4^+^ T-cell subset to the various available human metabolic reconstructions and pathway databases, we found that approximately 17% of the genes expressed in each CD4^+^ T-cell subset were found in the human metabolic reaction (HMR2) [30] database, whilst only ~5% of the genes were found in the Kyoto Encyclopedia of Genes and Genomes (KEGG) [31], and the Encyclopedia of Human Genes and Metabolism (HumanCyc) [32] pathway databases (**Figure S2**). The mapped genes for each CD4^+^ T-cell subset were assembled and listed as metabolic genes (MGs). A total of 638 MGs were common to both the human metabolic reconstructions and metabolic pathway databases, whilst 750 MGs were unique to HMR2 (**Figure 1B**). Since HMR2 had the highest coverage of MGs in our CD4^+^ T-cell datasets, it was used as a background model for the reconstruction of HTimmR.

When we investigated the differential expression of MGs between naïve (Thp), activated (Th0), and differentiated CD4^+^ T-cell subsets, we found that 853 MGs were differentially expressed (False Discovery Rate, FDR < 0.05), *i.e.*, up- or down-regulated in Th0 cells as compared to Thp cells (**Figure S2**). Similarly, 173, 506, 106 and 99 MGs were differentially expressed (FDR < 0.05) between Th1, Th2, Th17 and iTreg cells, respectively, as compared to Th0 cells at 72 hours of polarization (**Figure S2**). Gene ontology (GO) term mapping of biological processes linked to the MGs suggested that, 30.43% of MGs identified in CD4^+^ T-cell subsets encode metabolic processes (GO:0008152), whilst 31.52% and 1.09% encode cellular processes (GO:0009987) and immune responses (GO:0002376), respectively (**Figure 1C**).

### Reporter metabolites of specific lipids and amino acids are altered during activation of human CD4^+^ T-cells

Reporter metabolite (RM) analysis is an approach for the identification of metabolites in a metabolic network, around which significant transcriptional changes occur [33, 34]. RM analysis can predict hotspots in a metabolic network that are altered between two different conditions, in this case, Th0 *vs*. Thp cells.

The RM analysis suggests that biosynthetic intermediates of glycolysis and the tricarboxylic acid (TCA) cycle are altered upon CD4^+^ T-cell activation (**Figure 2A**). RMs such as acetyl-CoA (p=0.02), oxaloacetate (OAA) (p=0.01), itaconate (p=0.04) and itaconyl-CoA (p=0.04) were upregulated, whilst citrate (p=0.02) and fumarate (p=0.01) were downregulated in Th0 cells as compared to Thp cells. This implies that, upon activation of CD4^+^ T-cells, citrate can be diverted from the TCA cycle to form itaconate *via* aconitate. This phenomenon has been observed previously in macrophages [35], where itaconate is upregulated under inflammatory conditions to promote an anti-inflammatory response. However, it remains to be elucidated if itaconate accumulation during TCR activation plays a significant role in metabolic reprogramming [36] of T-cells.

**Figure 2.**
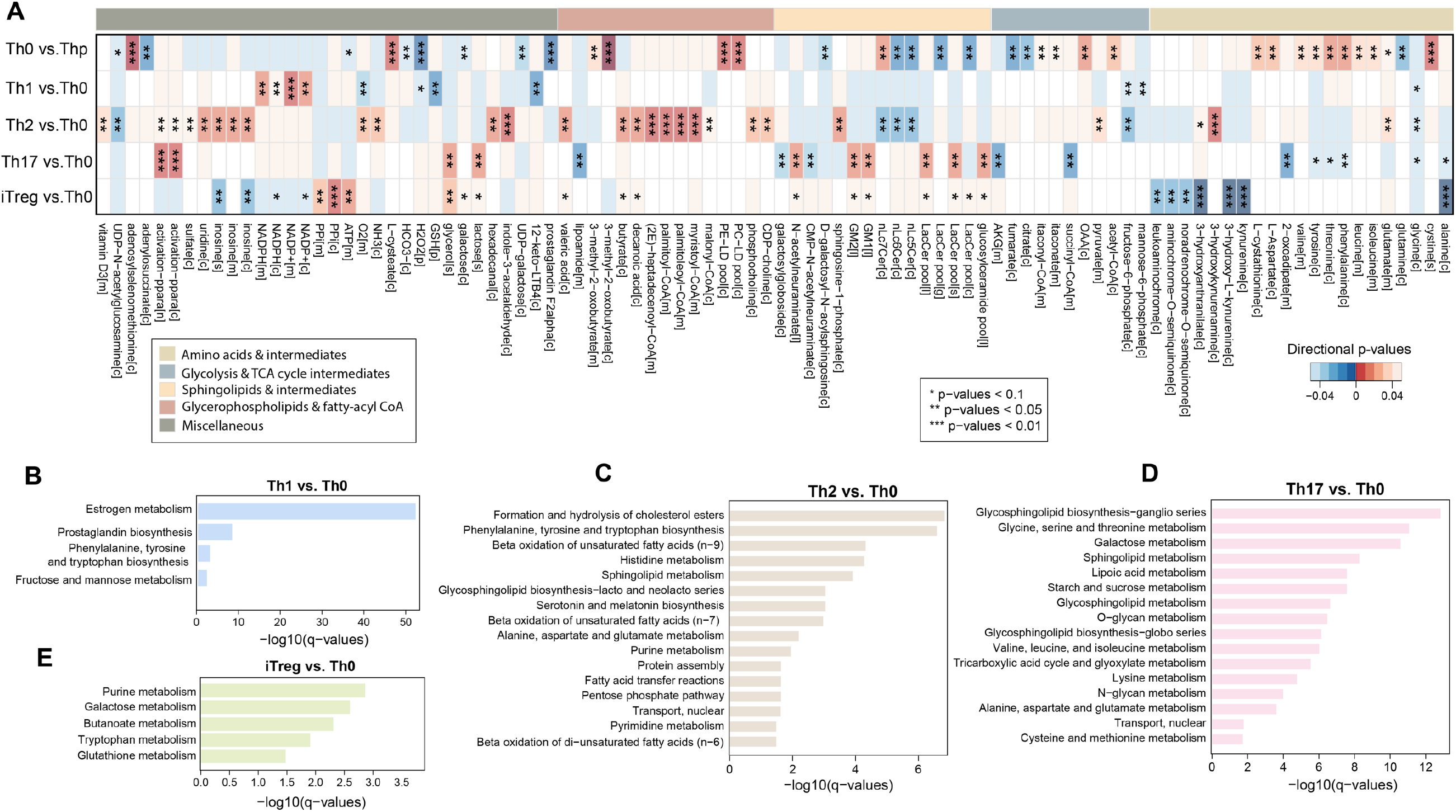
Reporter metabolites and overrepresented pathways of CD4^+^ T-cell activation and differentiation at 72 hours of polarization. (**A**) A heatmap of reporter metabolites (RMs) that are significantly (p < 0.05) up-(red color) or downregulated (blue color) or remained unchanged (white color), in the CD4^+^ T-cell subsets as compared to their paired control (Th0). The RMs were grouped by their metabolic subsystems / pathways and marked by the color bars. An asterisk (‘*’) denotes levels of statistical significance as determined by p-values. (**B-E**) Bar plots showing overrepresented (q < 0.05) reporter pathways (RPs) of the CD4^+^ T-cell subsets.

RMs of amino acids (valine (p=0.03), cystine (p=0.007), leucine (p=0.03), isoleucine (p=0.03), tyrosine (p=0.03), phenylalanine (p=0.01), and threonine (p=0.01)), and glycerophospholipids (phosphatidylcholine (PCs) (p=0.008) and phosphatidylethanolamine (PEs) (p=0.008)) were upregulated in Th0 as compared to Thp cells (**Figure 2A**).

Intriguingly, several intermediates of sphingolipids, particularly glycosphingolipid (GSL) pathways such as lactosylceramides (LacCers) (p=0.01) and D-galactosyl-N-acylsphingosine (p=0.03), were downregulated in Th0 as compared to Thp cells (**Figure 2A**).

Overrepresentation analysis of the RM pathways showed that, primarily, lipid (glycerophospholipids and GSL), and amino acid metabolism were altered (hypergeometric test, q-value < 0.05) upon CD4^+^ T-cell activation (**Figure S3**).

### Differentiation of human CD4^+^ T-cell subsets depicts regulation of unique metabolic pathways

RMs of Th1, Th2, Th17, and iTreg cells were identified at 72 hours of polarization (**Figure 2A**). In Th1 cells, we observed that RM pools of NADP+ (p=0.004) and NADPH (p=0.01) were upregulated as compared to Th0 cells. Differentiation of Th1 cells may alter intracellular levels of oxidative stress, as suggested by the downregulation of peroxisomal glutathione (GSH) (p = 0.01) and H_2_O_2_ (p = 0.009). Prostaglandins and leukotrienes (12-dehydro-leukotriene B4 and 12-oxo-leukotriene B3) were also downregulated (p = 0.017) (**Figure 2A**). Overrepresentation analysis of RM pathways suggests that prostaglandin biosynthesis, aromatic amino acid, estrogen, fructose and mannose metabolism are altered (hypergeometric test, q-value < 0.05) in the Th1 cells as compared to Th0 cells (**Figure 2B**).

In Th2 cells, mitochondrial fatty acyl-CoA, including myristoyl-CoA (p=0.002) and palmitoyl-CoA (p=0.005) RMs were upregulated as compared to Th0 cells. Intriguingly, these activated fatty acyl-CoAs might induce fatty-acid oxidation (FAO) and degradation. On the other hand, cytosolic malonyl-CoA (p=0.04) was elevated in Th2 cells. Malonyl-CoA inhibits the rate-limiting step of FAO, *i.e.*, transport of FAs to mitochondria *via* the carnitine shuttle. Our results suggest that there might be a trade-off between fatty acid synthesis (FAS) and FAO that underpins the functional differentiation of Th2 cells (**Figure 2A**). Several RM pathways such as β-oxidation of unsaturated fatty acids, alanine, aspartate, glutamate and histidine metabolism, the pentose phosphate pathway (PPP), GSL and nucleotide metabolism were altered (hypergeometric test, q-value < 0.05) in Th2 cells as compared to Th0 cells (**Figure 2C**).

RM analysis of Th17 showed a markedly-different pattern of regulation as compared to Th1 and Th2 cells. Several classes of GSLs (cerebrosides, gangliosides (GMs), N-acetylneuraminic acid (NANA)) were upregulated (p<0.05) in the differentiated CD4^+^ Th17 cells as compared to Th0 cells at 72 hours. In addition, some of the aromatic amino acids were downregulated (p<0.05) (**Figure 2A**). Overrepresentation analysis of RM pathways showed that sphingolipid, particularly GSL, amino acid metabolism, and other related metabolic processes were altered (hypergeometric test, q-value < 0.05) in Th17 cells as compared to Th0 cells (**Figure 2D**). Notably, GSLs (LacCers, GMs and NANA) were found to be a unique signature of differentiated CD4^+^ Th17 cells as compared to Th1 and Th2 cells. Some of these GSLs displayed a similar trend in the iTreg *vs.* Th0 cells, but the changes did not reach statistical significance (p=0.09). Additionally, RMs of the tryptophan / kynurenine pathways, *i.e.,* kynurenine, 3-hydroxyanthranilate, formylanthranilate and quinones, were downregulated (p=0.0008) (**Figure 2A**). Of note, tryptophan is metabolized to kynurenine by indoleamine 2,3-dioxygenase, an enzyme that is induced by pro-inflammatory cytokines [37]. In addition, several short and long chain fatty acids such as butyric (p=0.06), decanoic (p=0.06) and valeric (p=0.06) acids were upregulated in the iTreg *vs.* Th0 cells. Overrepresentation analysis of RM pathways showed that, tryptophan, glutathione, butanoate, galactose and purine metabolism were overrepresented in iTregs cells as compared to Th0 cells (**Figure 2E**).

### Dynamic regulation of molecular lipids in human CD4^+^ T-cell subsets

Taken together, RM analysis of CD4^+^ T-cell activation and differentiation predicted several classes of metabolites and pathways as being altered between the various subsets (**Figure 2** and **Figure S4**). RMs of lipids and amino acids were found to be the predominant classes significantly altered in CD4^+^ T-cells during activation and at 72 hours of differentiation. While the importance of amino acids in CD4^+^ T-cell differentiation is well-established [18, 38], the role of molecular lipids in the differentiation of human CD4^+^ T-cells remains uncharacterized. Furthermore, RMs of glycerophospholipids and GSLs were markedly altered in Th17 and iTreg cells as compared to Th1 and Th2 cells (**Figure 2**). Thus, the above findings directed us to investigate the dynamics of molecular lipids in CD4^+^ T-cell differentiation.

We profiled the molecular lipids of human CD4^+^ T-cell subsets (n=5) (**Figure S5−7**) using the established lipidomics platform, which is based on ultra-high performance liquid chromatography coupled to time-of-flight mass spectrometry (UHPLC-QTOFMS). Sparse Partial Least Square Discriminant Analysis (sPLS-DA) [39] of the lipidomic dataset reveled that the lipidome of resting naïve (Thp) cells was different from the activated or differentiated T-cell subsets (R^2^X = 0.933, R^2^Y = 0.988, N=7-fold cross-validated Q^2^ = 0.886; **Figure 3A**). Th17 and iTreg cells were different from each other, and also from the Th1 and Th2 cells (**Figure 3A**). Several classes of lipids, such as lysophosphatidylcholines (LPCs), PCs, PEs, sphingomyelins (SMs), ceramides (Cers) and triacylglycerols (TGs) were altered (regression coefficient, RC (>± 0.05) and Variable Importance in Projection (VIP) scores [40] > 1) in the CD4^+^ T-cell subsets at 72 hours of differentiation (**Figure 3B**).

**Figure 3.**
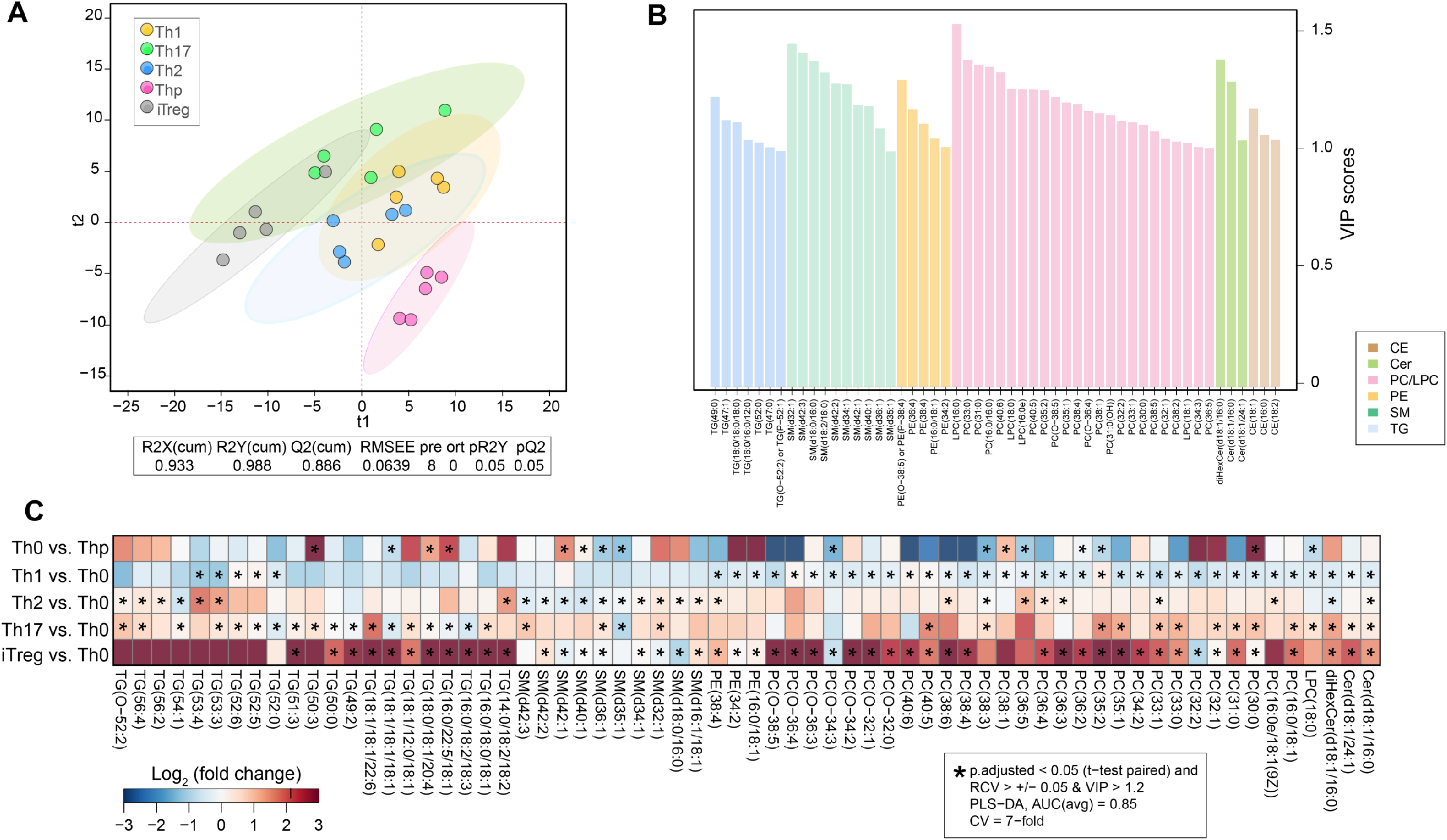
Lipidome of human CD4^+^ T-cell activation and differentiation. (**A**) Scatter / score plot for the PLS-DA classification model *(model performance: R^2^X = 0.933, R^2^Y = 0.988, N = 7-fold cross-validated (CV), Q^2^ = 0.886)*, showing differences in the lipidomes of the T-cell subsets, isolated from the umbilical cord of (n=5) healthy neonates. Ellipse denotes 95% confidence region. (**B**) A bar plot showing VIP scores of the lipids included in the PLS-DA classification model. The lipids are grouped and color-coded by their chemical classes. Different classes of lipids such as cholesterol esters (CEs), phosphatidylcholines (PCs), lysophosphatidylcholines (LPCs), phosphatidylethanolamines (PEs), sphingomyelins (SMs), ceramides (Cers), dihexosyl ceramides (diHexCers) and triacylglycerols (TGs) with (VIP scores >1) are shown. (**C**) Heatmap showing log_2_ fold changes (FC) of the significantly-altered lipids between T-cell subsets *versus* Th0 cells at 72 hours of polarization. Red color denotes increase whilst blue color denotes decrease, while white denotes no change. An asterisk (‘*’) denotes a significant difference in the levels of the lipids, as determined by the univariate (paired t-test, FDR < 0.05) and multivariate (PLS-DA; abs(RCV) > 0.05 and VIP > 1.2) analyses.

We combined multivariate (sPLS-DA, n=7-fold cross validation, CV) and univariate (paired t-test) approaches to identify cellular lipid signatures that were altered significantly (area under the curve (AUC) ~0.85, regression coefficient RC (>± 0.05) and VIP scores > 1.2 and paired t-test, p-adjusted < 0.05) between the CD4^+^ T-cell subsets, and their paired Th0 (controls) (**Figure 3C**). We found that PCs, LPCs, SMs and Cers were altered in the T-cell subsets during activation and early differentiation (**Figure 3C)**. The majority of cellular PCs were upregulated in the Th2, Th17 and iTreg cells, whilst these were downregulated in Th1 cells, as compared to their paired Th0 cells (**Figures 3C** and **S8**). Cer(d18:1/16:0), Cer(d18:1/24:1) and diHexCer(d18:1/16:0) were elevated in Th17 and iTreg cells. Mostly, SMs were altered in Th2 and iTreg cells. Some SMs (SM(d32:1), SM(d36:1) and SM(d42:1)) were upregulated in Th17 cells (*vs.* Th0), whilst downregulated in iTreg cells (*vs.* Th0) (**Figures 3C** and **S9**). Cellular TGs were markedly elevated in iTreg and Th17 cells (**Figures 3C** and **S10**). The lipidome data suggests that glycerophospholipids (PCs, PEs and LPCs) and sphingolipids (Cers, GSLs and SMs) are the major indicators of CD4^+^ T-cell differentiation (**Figure 3B-C**).

### Dynamic regulation of ceramides and glycosphingolipids during the early differentiation of human CD4^+^ Th17 and iTreg cells

GSLs and Cers play an important role in maintaining the integrity of the plasma membrane. They are involved in cellular signaling, proliferation, endocytosis, and modulate cellular responses to inflammatory and apoptotic stress signals [41, 42]. However, the functional role of these metabolites in CD4^+^ T-cell activation and differentiation remains unknown [41]. Here, RM predictions and lipidome analysis of human CD4^+^ T-cells have, together, identified several species of GSLs that were altered in Th17 (*vs*. Th0) and iTreg cells (*vs.* Th0 cells) at 72 hours of differentiation (**Figures 2A** and **3B-C**).

Next, by applying RM analysis, we investigated the regulation of GSLs and Cers in Th17 and iTreg cells during the first 48 hours of polarization. RM analysis was performed between Th17 *vs*. Th0 cells at 0.5, 1, 2, 4, 6, 12, 24, 48 hours of differentiation. Initially, the RM of Cers were elevated at 1 hour, and subsequently, there was a transient decrease at 2 hours, followed by an increase at 12, 24, 48 hours of differentiation (**Figure S11**). A similar pattern of regulation was also observed in ceramide 1-phosphate (C1P), an active intermediate of sphingolipid metabolism (**Figure S11**).

In iTreg cells, several RMs of the sphingolipid pathway such as digalactosylceramides, D-galactosyl-N-acylsphingosine, UDP-galactose and diHexCers, particularly, lactosylceramides (LacCers) were upregulated (*vs.* Th0) by 6 hours of polarization (**Figure S12**). However, no change in the levels of LacCers were observed in Th17 cells at these early time-points (**Figure S11**).

### Targeted lipid measurements reveal the regulation of ceramide levels in human CD4^+^ Th17 and iTreg cells at 72 hours of differentiation

A targeted lipidomics experiment was designed to measure the levels of Cers and GSLs (*i.e.* hexosylceramides (HexCers) and diHexCers in human CD4^+^ Th17 (n=3) (**Figure S7**) and iTreg cells (n=5) (**Figure S6**), at 72 hours of differentiation (**Figure 4** and **Table S1**). Increased levels of Cers, and decreased levels of HexCers, were found in both Th17 and iTreg cells, except for HexCer(18:1/24:0), which was elevated (p=0.04) in iTreg cells (**Figure 4H**). Intriguingly, diHexCers (d18:1/16:0) were elevated in Th17 *vs.* Th0 cells (p=0.001), while these changes were not apparent (p>0.05) in iTreg cells *vs.* Th0 cells (**Figure 4I**).

**Figure 4.**
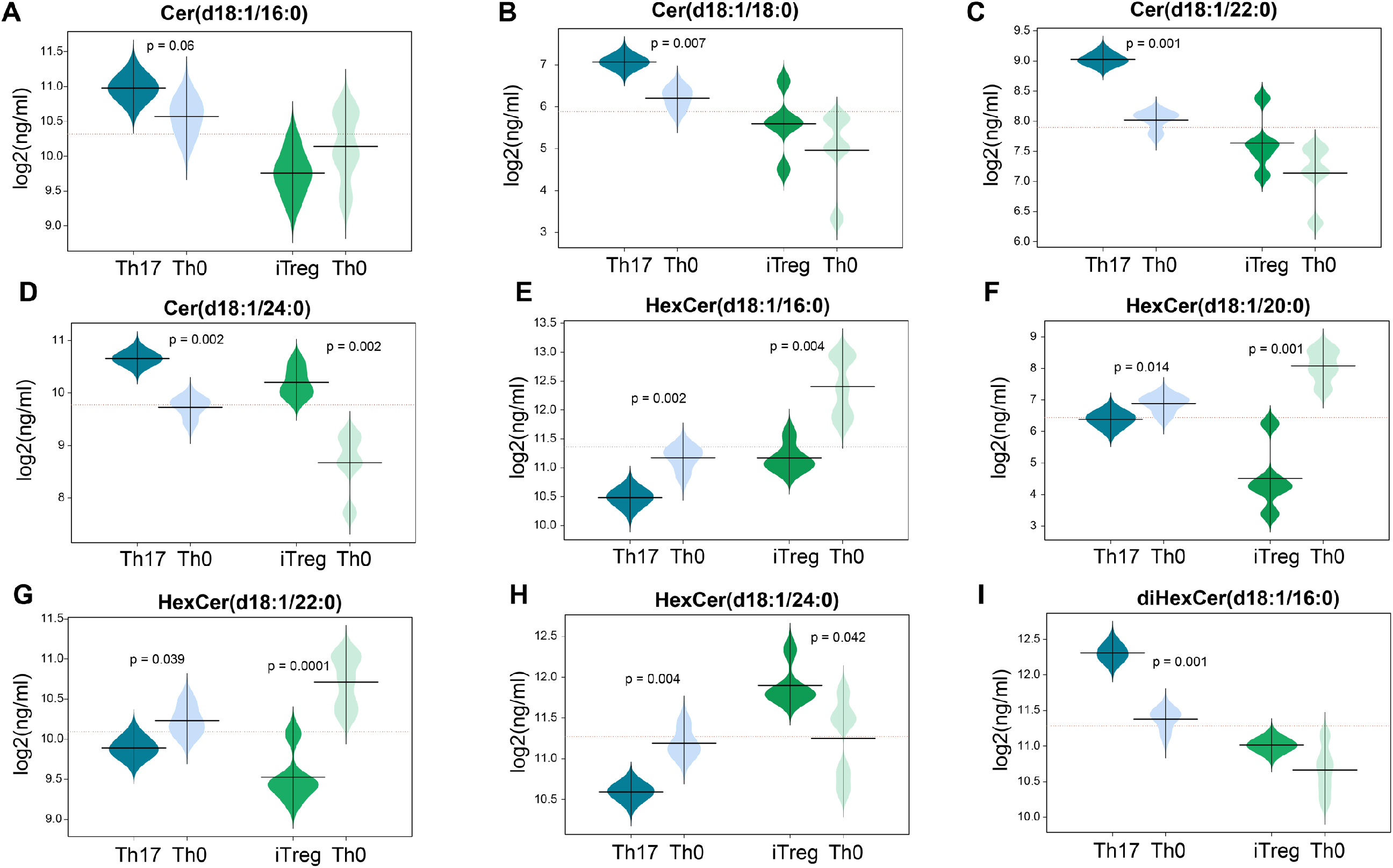
Targeted quantification of ceramide levels in Th17 and iTreg cells. (**A-D**) Beanplots showing ceramide levels (log_2_(ng/mL)) measured in Th17 and iTreg cells isolated from the umbilical cord of (n=5) healthy neonates, and their paired controls (Th0) at 72 hours of polarization. (**E-I**) Showing the cellular levels of HexCers and diHexCers in the Th17 and iTreg cells along with their paired controls (Th0), at 72 hours of polarization. Significant differences (paired t-test, p < 0.05) are shown by the p-values. The dotted line denotes the mean of the population, and the black dashed lines in the bean plots represent the group mean.

There was congruence between the results obtained from GSMM-RM predictions (**Figure 2A**), non-targeted (**Figure 3B-C**) and targeted (**Figure 4**) lipidomics measurements showing that GSLs (specifically HexCers and diHexCers) were elevated in CD4^+^ Th17 cells (p<0.05).

### Relative contribution of the sphingolipid pathways to the production of ceramides in Th17 cells

We evaluated the effect of various sphingolipid pathways on Cer production in Th17 cells at 72 hours of differentiation (**Figure 5A**). An *in-silico* knockout (KO) of the Th17-cell-specific pathways was simulated using GSMM, where each sphingolipid pathway was knocked out iteratively, one at a time, and the percentage of maximum flux contributions for Cer production *via* eight different metabolic reactions were estimated (**Figure 5A-B**). Optimization of Cer production in a wild type (WT) model suggests that, ~40% of the total flux of Cer production can be carried by *de novo* synthesis, *i.e.*, by conversion of dihydroceramide to Cer (**Figure 5A-B**). As expected, KO of serine palmitoyltransferase (SPT) pathway, a rate-limiting-step in the *de novo* sphingolipid synthesis pathway [43], decreased the total flux of Cer production, however, knockout could not completely abolish Cer formation (**Figure 5B**). This is due to the presence of redundant sphingolipid pathways which are able to replenish Cers in Th17 cells (**Figure 5A-B**).

**Figure 5.**
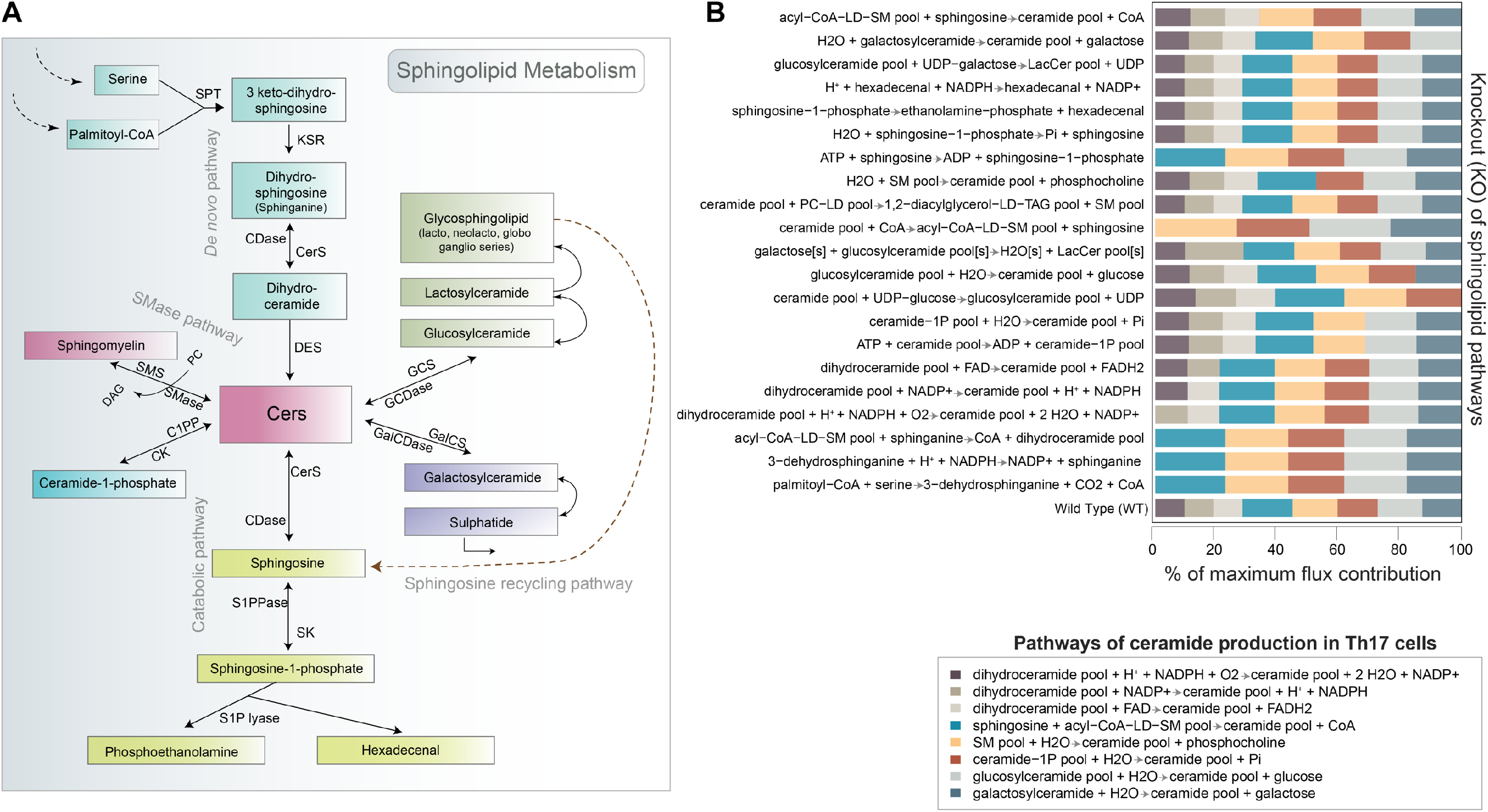
Regulation of sphingolipid pathways in human CD4^+^ T-cells. (**A**) An Illustration of sphingolipid metabolism in human CD4^+^ T-cells. (**B**) *In silico* knockout (KO) analysis showing, the % of maximum flux contribution of different sphingolipid pathways towards the Cer production, by knocking out (one-by-one) an alternate sphingolipid pathway. Abbreviations: C1P, ceramide 1-phosphate; CDase, ceramidase; CerS, ceramide synthase; DES, dihydroceramide desaturase; GalCDase, galactosidase; GCDase, glucosidase; S1P, sphingosine 1-phosphate; S1PPase, sphingosine phosphate phosphatases; SK, sphingosine kinase; SMase, sphingomyelinase pathway; SPT, serine palmitoyl-CoA transferase.

### Effects of *SPTLC123* silencing on ceramide levels in human CD4^+^ Th17 cells

*De novo* biosynthesis of Cer starts with the *serine palmitoyl transferase* (*SPT*) pathway, a rate-limiting step that aids in the condensation of serine and palmitoyl-CoA by an enzyme complex called serine palmitoyl transferase (*SPTLC*) (**Figures 5A** and **6A**). Currently, there are three major *SPTLC* subunits identified, including *SPTLC1, SPTLC2* and *SPTLC3*, which are known to be expressed in humans [44].

Based on our RNA-Seq data, the three *SPTLC* subunits are expressed during human Th17 cell differentiation. Among these, *SPTLC1* and *SPTLC2* showed higher levels of expression than *SPTLC3* (**Figure S13**). To examine changes in Cer biosynthesis during Th17 cell development, we simultaneously silenced these three SPTLC subunits (*SPTLC123* triple knockdown, TKD) using small interfering RNAs (*siRNAs)*. As illustrated in (**Figure 6B-E**), the three *siRNAs* successfully downregulated their targets. Importantly, silencing of *SPTLC* decreased the expression of the proinflammatory cytokines IL17A (p=0.08) and IL17F (p=0.001) in Th17 cells at 72 hours of following cell activation (**Figure 6F-G**), suggesting that Cer synthesis is vital for Th17 cell function.

**Figure 6.**
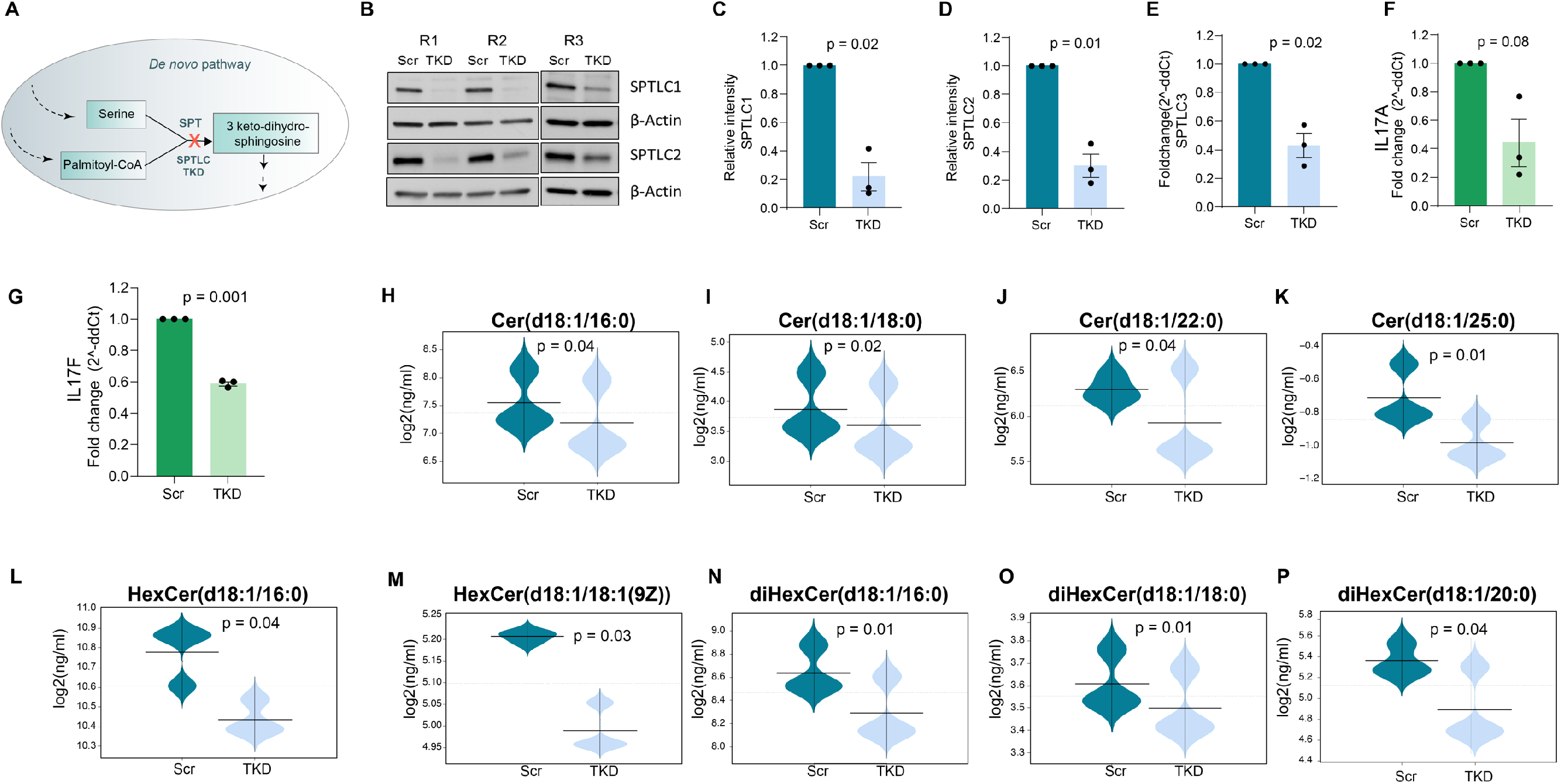
Effect of *SPTLC* deficiency on serine palmitoyltransferase (SPT) *de novo* pathway and Th17 differentiation. (**A**) Illustration of *SPT* (*SPTLCs*) TKD. (**B-E**) Immunoblots and corresponding quantified intensities of *SPTLC1* and SPTLC2 protein expression at 24 hours and fold changes of *SPTLC3* mRNA expression by quantitative real-time PCR at 72 hours upon *SPTLC* TKD in Th17 cell differentiation (Scr vs *SPTLC* TKD; n=3; paired t-test, p<0.05). (**F-G**) Fold changes of IL17A and IL17F mRNA expression (Scr *vs. SPTLC* TKD) at 72 hours of Th17 cell differentiation (n=3; paired t-test, p<0.05). H-P) Beanplots showing the targeted quantification levels of (log_2_ (ng/mL)) Cers, HexCers, and diHexCers measured in control (Scr) and *SPTLC*-deficient Th17 cells at 72 hours (n=3). Significant differences (paired t-test, p < 0.05) are shown by the p-values. The dotted line denotes the mean of the population, and the black dashed lines in the bean plots represent the group mean.

Similarly, through a mass spectrometry-based targeted lipidomics experiment, we measured the Cer levels in these Th17 (TKD) cells at 72 hours of polarization. Several species of Cers and GSLs (HexCers, diHexCers) were significantly decreased in *SPTLC*-deficient Th17 cells (**Figure 6H-P**). Most importantly, diHexCers levels were significantly reduced upon *SPTLC* silencing, suggesting that regulation of diHexCers is associated with the Th17 differentiation and effector function (**Figure 6N-P**). Overall, these results suggest that silencing of the three *SPTLC* subunits negatively influences the development and secretory function of human CD4^+^ Th17 cells (**Figure 6F-G**).

### *Effect of UGCG silencing on* hexosyl- and lactosylceramide synthesis in human CD4^+^ Th17 cells

diHexCers (LacCers) can be generated from HexCers (GlcCers) which, in turn, are produced from Cers and glucose, catalyzed by *glucosylceramide synthase (GCS)*(EC 2.4.1.80), encoded by the *UGCG* gene. This is the first committed step in the production of GlcCer-related GSLs [45] (**Figure 5A** and **Figure 7A,D**). Results from the *SPTLC* gene silencing experiment in Th17 cells showed an effective downregulation of diHexCers (**Figure 6N-P**). Further, these findings guided us to evaluate the importance of the GCS-pathway for the production of GlcCers and diHexCers in human CD4^+^ Th17 cells.

**Figure 7.**
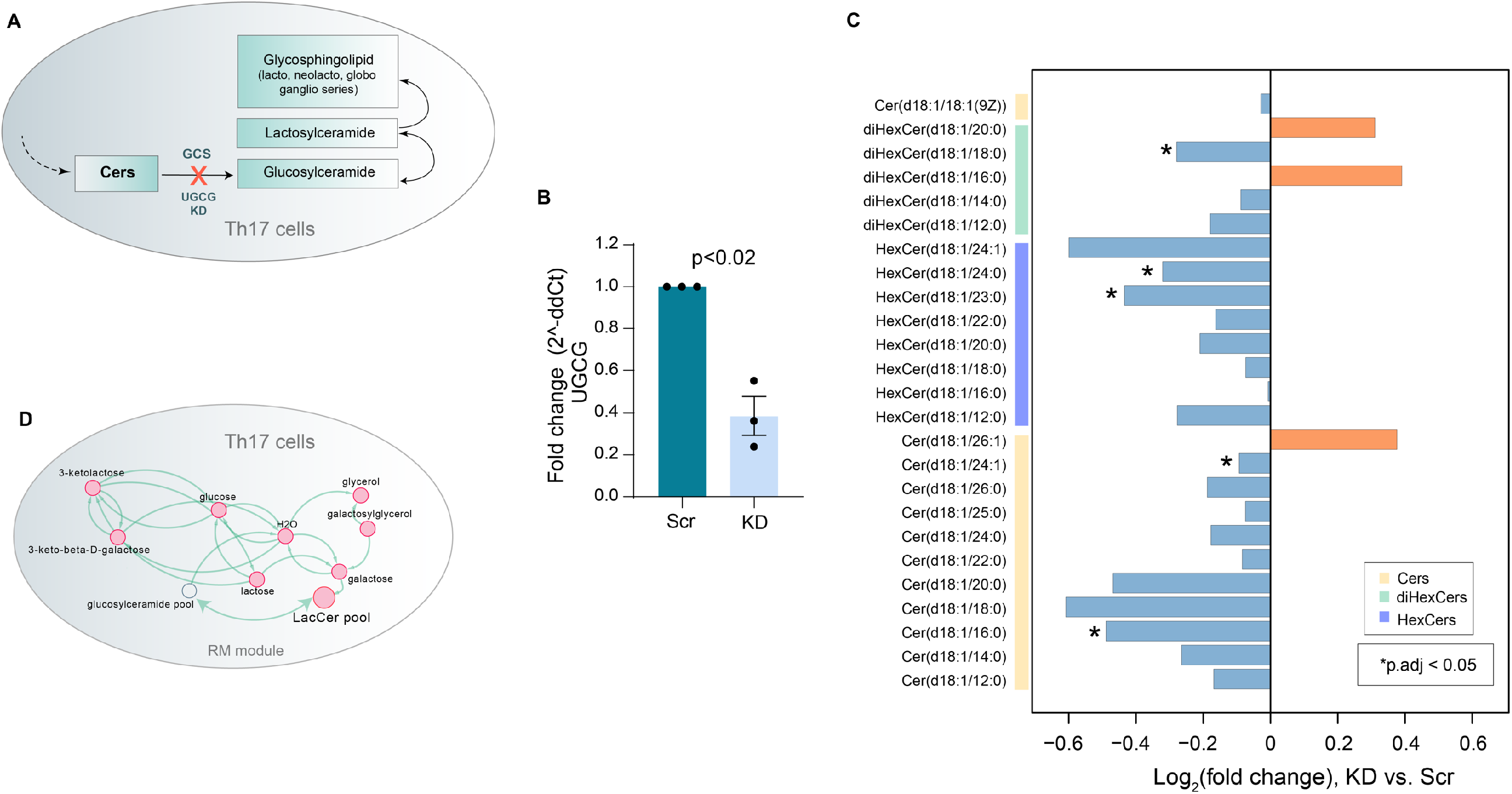
Targeted quantification of the Cer and GSL levels in the *UGCG*-deficient Th17 cells. (**A**) Illustration of GCS pathway and (*UGCG*) knockdown. (**B**) Fold change of *UGCG* gene expression in control (Scr) and *UGCG*-deficient Th17 cells at 12 hours (n=3; paired t-test, p<0.05). (**C**) Bar plot showing log_2_ fold changes of the Cers, HexCers, and diHexCers measured in *UGCG*-deficient (KD) *vs.* control (Scr) Th17 cells at 72 hours (n=3). (**D**) Elevated RM pathway module (p < 0.05) comprising GlcCers, diHexCers, and their congeners in Th17 cells, identified in this study.

Similarly, by taking a *siRNA*-mediated silencing approach, we knocked down expression of the *UGCG* gene (**Figure 7B**), which encodes GCS in Th17 cells (**Figure S13**). Although expression of the IL17 cytokine was not influenced by *UGCG* knockdown in Th17 cells, we decided to determine changes in the HexCers, diHexCers and SMs production in *UGCG*-deficient Th17 cells. As expected, several species of Cers, HexCers (GlcCers and / or GalCers) and diHexCers (except diHexCer(d18:1/16:0)) were decreased in the *UGCG*-silenced Th17 cells (**Figure 7C**). A decrease in Cer levels implies that Cer can be diverted to other sphingolipid pathways (*CerS*, *SMS*, *C1PP*; **Figure 5A**), and thus might enhance the production of sphingosine, SMs and ceramide-1-phosphate, respectively, which, in turn, regulate cytokine production. Intriguingly, several species of SMs were elevated in the *UGCG*-silenced Th17 cells suggesting that, production of SMs *via* the SMS-pathway was enhanced with a pertinent decrease in Cers and GSLs (**Figure 5A** and **Figure S14**).

## DISCUSSION

We showed that human CD4^+^ T-cell subsets*, i.e.*, Th1, Th2, Th17 and iTreg cells, undergo both common and subset-specific metabolic alterations, in order to both successfully differentiate and subsequently carry out their specific functions. RM analysis suggests that, upon the activation of naïve T-cells, the levels of the amino acids increase, while the majority of these amino acids then decrease during towards a Treg phenotype. Indeed, iTreg cells are thought to be less dependent on amino acids [46]. For instance, depletion of glutamine can skew the differentiation of CD4^+^ T-cells towards a Treg lineage [47].

Cers are key intermediates of sphingolipid metabolism, composed of sphingosine base and a fatty acyl chain (C14:0 - C26:0) [48]. Cers are important for T-cell activation and differentiation at multiple levels, such as intracellular signal transduction, modulation of membrane fluidity, receptor clustering and by contributing to CD95-mediated cell death *via* multiple mechanisms [49]. However, the Cer pathways are highly redundant. One of the key, novel observations from our RM and lipidome analyses was that several species of Cers and GSLs are markedly altered in Th17 and iTreg cells at 72 hours of differentiation. The observed alterations in the levels of Cers and GSLs (HexCers, diHexCers) were prominent in the Th17 cells as compared to the iTreg cells. diHexCers (LacCers) were markedly-high in the differentiated Th17 *vs.* Th0 cells. Furthermore, *in vitro* KD experiments substantiated the essentiality of sphingolipid metabolic pathways *(SPT, GCS)* in the formation of Cers and GlcCers / diHexCers, and these are intrinsically linked to proinflammatory cytokine (IL17A and IL17F) expression in Th17 cells. Several species of diHexCers were decreased in the knockdowns, suggesting that accumulation of diHexCers is required for Th17 differentiation. In addition, RM analysis of Th17 cells showed a persistent increase in the Cer pool from 12 hours until 72 hours of polarization.

An earlier study observed accumulation of Cers in Treg cells as a consequence of low sphingomyelin synthase *SMS1* (encoded by *Sgms1*), an enzyme catalyzing the conversion of Cers and PCs to diacylglycerols and SMs [42]. In line with this, several species of SMs measured in our study were shown to be decreased in the iTregs *vs.* Th0, and increased in Th17 cells *vs.* Th0, while few SMs (SM(d16:1/18:1) and SM(d42:3)) elevated in both iTregs and Th17 cells (**Figure 3C**). In Treg cells, FOXP3 directly binds to *Sgms1* to suppress *SMS1*, and retroviral overexpression of FOXP3 in Jurkat human T lymphocytes decreased the expression of *SGMS1* [42, 50]. The accumulation of Cers constrains *SET* activity towards protein phosphatase A (*PP2A*). Intriguingly, *PP2A* can suppress mTORC1 activity and promotes Treg and Th17 cell differentiation [42, 51, 52]

Furthermore, several studies indicate that co-expression of CD39 (*ENTPD1*) and CD161 (*KLRB*) in Th17 cells increases the activity of acid sphingomyelinase (*ASM*), an enzyme encoded by the gene *SMPD1*, which hydrolyzes SMs to form Cers and phosphorylcholine, in turn leading to an increase in the Cer pool [53–55]. Although *ENTPD1* levels remain unchanged during the early differentiation of iTreg and Th17 cells; *KLRB* is specifically downregulated in Th17 cells [56, 57]. Further, *SMPD1* is upregulated in iTreg cells [56], suggesting intrinsic regulation of Cers in both Th17 and iTreg cells.

Taken together, our study identified several common and subset-specific metabolic signatures and pathways in human CD4^+^ T-cells, for their activation, and functional differentiation. This enabled us to improve understanding of how molecular lipids are regulated in different subsets of CD4^+^ T-cells. We demonstrated the essentiality of ceramide and glycosphingolipid synthesis pathways for Th17 differentiation and effector function. Our study may, therefore, provide a comprehensive resource for identifying CD4^+^ T-cell-specific metabolic pathways and useful targets for their selective manipulation under disease conditions characterized by an imbalance of Treg / Th17 cells [5, 10, 24, 29].

Our study may also offer clues about the poorly-understood overlap in co-morbidities between these immune-mediated diseases and metabolic diseases, such as has been found to occur in COVID19 [58]. Obesity is commonly associated with chronic elevation of circulating fatty acids, which results in the accumulation of, among others, toxic lipids, such as Cers, in peripheral cells / tissues – a phenomenon referred to as lipotoxicity [59]. The presence of elevated levels of ceramides in conditions such as obesity and insulin resistance may thus skew the Treg / Th17 balance towards the pro-inflammatory Th17 phenotype, which is also being reported as one of the notable hallmarks of severe COVID19 [60, 61].

## METHODS

### Human CD4^+^ T-cell Isolation, activation, and differentiation

CD4^+^ T-cells were isolated from human umbilical cord blood as described previously [56, 62, 63]. For Th17 cell differentiation, isolated CD4^+^ cells were activated with a combination of plate-bound anti-CD3 (750 ng/24-well culture plate well; Immunotech/Beckman Coulter REF # IM-1304) and soluble anti-CD28 ((1ug/mL; Immunotech/Beckman coulter REF # IM1376) antibodies in serum-free X-Vivo 20 medium (Lonza), in the absence (Th0) or presence (Th17) of IL-6 (20ng/ml, Roche, Cat# 11138600 001); IL-1β (10ng/ml, R&D Systems Cat # 201 LB); TGF-β1 (10ng/ml, R&D Systems Cat# 240); anti-IL-4 (1 μg/ml) R&D Systems Cat# MAB204) and anti-IFN-γ (1 μg/ml R&D Systems Cat#MAB-285). Differentiation of Th17 cells was confirmed by measuring IL-17 expression by quantitative real-time PCR, at 72 hours of Th17 / Th0 culturing [62].

For iTreg cell culturing, after of CD25+ cells, done using LD columns and a CD25 depletion kit (Miltenyi Biotec), CD4^+^CD25^−^ cells were activated with plate-bound anti-CD3 (500 ng/24-well culture plate well) and soluble anti-CD28 (500 ng/mL) at a density of 2 × 10^6^ cells/mL of X-vivo 15 serum-free medium (Lonza). For iTreg differentiation, the medium was supplemented with IL-2 (12 ng/mL), TGF-β (10 ng/mL) (both from R&D Systems), all-trans retinoic acid (ATRA) (10 nM; Sigma-Aldrich), and human serum (10%) and cultured at 37°C in 5% CO_2_. Control Th0 cells were stimulated with plate-bound anti-CD3 soluble anti-CD28 antibodies without cytokines. For confirmation of iTreg cell differentiation, we used intracellular staining to measure, at 72 hours of iTreg culturing, expression of FOXP3 which is the major transcription factor driving Treg differentiation. Intracellular staining was performed using buffer sets of Human Regulatory T-cell Staining Kit (eBioscience/Thermo Fisher Scientific), following the manufacturer’s protocol. The following antibodies were used: anti-human FOXP3-PE (eBioscience, Cat. No. 12-4776-42) and rat IgG2a isotype control (eBioscience, Cat. No. 72-4321-77A). All samples were acquired by a flow cytometer (LSRII) and analyzed either with FlowJo (FLOWJO, LLC) or with Flowing Software [56].

Th1 and Th2 cell differentiation were done as described previously [64]. Briefly, purified naive CD4^+^ T-cells were activated with plate-bound anti-CD3 (500 ng/24-well culture plate well) and 500 ng/ml soluble anti-CD28 and cultured in the absence (Th0) or presence of 2.5 ng/ml IL-12 (R&D Systems) (Th1) or 10 ng/ml IL-4 (R&D Systems) (for Th2). At 48 hours following the activation of the cells, 17 ng/ml IL-2 (R&D Systems) was added to the cultures. Differentiation of Th1 and Th2 cells was confirmed by measuring (using flow cytometry) the expression of T-bet and Gata3 at 72 hours after cell activation. Briefly, cells were fixed and permeabilized using the Intracellular Fixation & Permeabilization Buffer Set (eBioscience / Thermo Fisher Scientific), according the manufacturer’s protocol. The following antibodies were used: anti-human GATA3-PE (eBioscience, 12-9966), anti-human T-bet-BV711 (BD, 563320) and corresponding isotype controls (BV711 Mouse IgG1, BD, 563044 and PE Rat IgG2b, eBioscience, 12-4031-82). Samples were acquired by BD LSRFortessa™ cell analyzer and data were analyzed using FlowJo software (FLOWJO, LLC).

### siRNA mediated gene knockdown

For *SPTLC* triple knock down (TKD) and *UGCG* single knock down (KD) experiments, freshly-isolated CD4^+^ cells were suspended in Optimem I (Invitrogen) and transfected with siGenome SMARTpool small interference RNA (siRNA) oligonucleotides (Dharmacon) using the nucleofection technique by Lonza. Scrambled non-targeting siRNA (5’-AAUUCUCCGAACGUGUCACGU-3’) was used as control (Sigma). Briefly, four million cells were transfected with 12 μg of *SPTLC*-targeting siRNAs (4 μg of SMARTpool *SPTLC1* siRNA M-006673-02; 4 μg of SMARTpool *SPTLC2* siRNA M-006674-01; and 4 μg of SMARTpool *SPTLC3* siRNA M-010285-02) or 12 μg of Scramble siRNA. For UGCG single knockdown experiments 12 μg of UGCG-targeting siRNA (siGenome SMARTpool, M-006441-02) were used. Cells were rested for 24h in RPMI 1640 medium (Sigma-Aldrich) supplemented with penicillin/streptomycin, 2 mM L-glutamine and 10% FCS and subsequently activated and cultured under Th17 conditions. *SPTLC1* and *SPTLC2* knockdown was validated by western blot at 24 hours, *UGCG* and *SPTLC3* knockdown was determined using quantitative real-time PCR (at 12 and 72 hours, respectively).

### Western blot

Fresh cell samples were lysed in RIPA buffer (Thermo) supplemented with complete EDTA-free Protease inhibitor cocktail and phosphatase inhibitors (Roche) and sonicated on a Bioruptor (Diagenode). Protein concentration was determined using DC Protein assay (Biorad). After boiling in 6× loading dye (330 mM Tris-HCl, pH 6.8; 330 mM SDS; 6% β-ME; 170 μM bromophenol blue; 30% glycerol), the samples were loaded on Mini-PROTEAN TGXPrecast Protein Gels (BioRad Laboratories) and transferred to PVDF membranes (Trans-Blot TurboTransfer Packs, BioRad Laboratories). The following primary antibodies were used: *SPTLC1* (sc-374143, Santa Cruz), *SPTLC2* (ab236900, abcam) and beta-actin (A5441, Sigma-Aldrich).

### TaqMan Quantitative Real-time PCR

Total RNA was extracted using the AllPrep DNA/RNA/miRNA Universal Kit (QIAGEN) and treated in-column with DNase (RNase-Free Dnase Set; QIAGEN) for 15 minutes. For quantitative real-time PCR purified RNA was treated with DNase I (Invitrogen) to ensure complete removal of genomic DNA followed by cDNA synthesis with SuperScript II Reverse Transcriptase (Invitrogen). Quantitative real-time PCR (qPCR) was performed using the TaqMan® Gene Expression *UGCG* Assay ID:Hs00916612_m1 and SPTLC3 Assay ID:Hs00217867_m1 (Thermo Fisher Scientific) or KAPA™ probe fast qPCR Master Mix (Kapa Biosystems) and Universal ProbeLibrary probes (Roche Applied Science) with custom ordered primers. The qPCR runs were analyzed with Applied Biosystems QuantStudio 12K Flex Real-Time PCR System. All reactions were performed in triplicate.

### Analysis of molecular lipids

The samples were randomized and extracted using a modified version of the previously-published Folch procedure [65]. Briefly, 150 μL of 0.9% NaCl was added to cell pellets, and samples then vortexed and ultrasonicated for 3 minutes. Next, 20 μL of the cell suspension was mixed with 150 μL of the 2.5 μg mL^−1^ internal standards solution in ice-cold CHCl3:MeOH (2:1, v/v). The internal standard solution contained the following compounds: 1,2-diheptadecanoyl-sn-glycero-3-phosphoethanolamine (PE (17:0/17:0)), N-heptadecanoyl-D-erythro-sphingosylphosphorylcholine (SM(d18:1/17:0)), N-heptadecanoyl-D-erythro-sphingosine (Cer(d18:1/17:0)), 1,2-diheptadeca-noyl-sn-glycero-3-phosphocholine (PC(17:0/17:0)), 1-heptadecanoyl-2-hydroxy-sn-glycero-3-phosphocholine (LPC(17:0)) and 1-palmitoyl-d31-2-oleoyl-sn-glycero-3-phosphocholine (PC(16:0/d31/18:1)). These were purchased from Avanti Polar Lipids, Inc. (Alabaster, AL, USA). In addition, triheptadecanoin (TG(17:0/17:0/17:0)) was purchased from (Larodan AB, (Solna, Sweden). The samples were vortexed and incubated on ice for 30 min after which they were centrifuged at 7800 × g for 5 min. Finally, 60 μL from the lower layer of each sample was collected and mixed with 60 μL of ice cold CHCl3:MeOH (2:1, *v/v*) in LC vial. The total protein content in cells was measured by the Bradford method [66].

The UHPLC-QTOFMS analyses were done in a similar manner to as described earlier, with some modifications [67, 68] on two separate instruments. The initial lipidomic results were acquired on a UHPLC-QTOFMS system from Agilent Technologies (Santa Clara, CA, USA) combining a 1290 Infinity LC system and 6545 quadrupole time of flight mass spectrometer (QTOFMS), interfaced with a dual jet stream electrospray (dual ESI) ion source. MassHunter B.06.01 software (Agilent Technologies, Santa Clara, CA, USA) was used for all data acquisition. The SM results for *UGCG*-silenced Th17 cells data was acquired on a UHPLC-QTOF system from Bruker (Bruker, Billerica, MA, USA) combining an Elute UHPLC binary pump and an Impact II system QTOF system. The samples for this experiments were the same extracts that the Cer data was acquired from and had SM(18:1/17:0) spiked in prior to acquisition. The data was acquired using the Hystar suite of software. MZmine 2 was used for all the untargeted data processing [69].

Chromatographic separation was performed using an Acquity UPLC BEH C18 column (100 mm × 2.1 mm *i.e.*, 1.7 μm particle size) and protected using a C18 precolumn, both from Waters Corporation (Wexford, Ireland). The mobile phases were water (phase A) and acetonitrile:2-propanol (1:1, *v/v*) (phase B), both containing 1% 1M ammonium acetate and 0.1% (*v/v*) formic acid ammonium acetate as ionization agents. The LC pump was programmed at a flow rate of 0.4 mL min^‒1^ and the elution gradient was as follows: from min 0–2, the percentage of phase B was modified from 35% to 80%, from min 2-7, the percentage of phase B was modified from 80% to 100% and this final percentage held for 7 min. A post-time of 7 min was used to regain the initial conditions for the next analysis. Thus, the total analysis time per sample was 21 min (including postprocessing). The settings of the dual ESI ionization source were as follows: capillary voltage 3.6 kV, nozzle voltage 1500 V, N_2_ pressure in the nebulizer 21 psi, N_2_ flow rate and temperature as heat gas 11 L min^−1^ and 379 °C, respectively. Accurate mass spectra in MS scan were acquired in the m/z range 100 – 1700 in positive ion mode.

MS data were processed using the open source software MZmine 2.53 [70]. The following data processing steps were applied to the raw MS data: (1) Crop filtering with a m/z range of 350 **–** 1200 m/z and a retention time (RT) range of 2 to 15 minutes; (2) Mass detection with a noise level of 900; (3) Chromatogram builder with a min time span of 0.08 minutes, minimum height of 900 and m/z tolerance of 0.006 m/z or 10.0 ppm; (4) Chromatogram deconvolution using the local minimum search algorithm with a 70% chromatographic threshold, 0.05 min minimum RT range, 5% minimum relative height, 1200 minimum absolute height, a minimum ration of peak top/edge of 1.2 and a peak duration range of 0.08 - 1.01 minutes; (5) Isotopic peak grouper with a m/z tolerance of 5.0 ppm, RT tolerance of 0.05 minute, maximum charge of 2 and with the most intense isotope set as the representative isotope; (6) Join aligner with m/z tolerance of 0.009 or 10.0 ppm and a weight of 2, RT tolerance of 0.1 minute and a weight of 1 and with no requirement of charge state or ID and no comparison of isotope pattern; (7) Peak list row filter with a minimum of 7 peaks in a row (10% of the samples); (8) Gap filling using the same RT and m/z range gap filler algorithm with an m/z tolerance of 0.009 m/z or 11.0 ppm; (9) Identification of lipids using a custom database (based on UHPLC-MS/MS data using the same lipidomics protocol, with RT data and MS and MS/MS) search with an m/z tolerance of 0.009 m/z or 10.0 ppm and a RT tolerance of 0.2 min. In general, lipids were identified at the total number of carbons and double bonds in the structure as there was insufficient evidence to assign the specific acyl chains. Where the acyl chains are identified these have been confirmed with MS/MS level experiments and/or authentic standards. (10) Normalization using internal standards (PE (17:0/17:0), SM (d18:1/17:0), Cer (d18:1/17:0), LPC (17:0), TG (17:0/17:0/17:0) and PC (16:0/d30/18:1)) for identified lipids and closest internal standard (based on RT) for the unknown lipids, followed by calculation of the concentrations based on lipid-class calibration curves.

Identification of lipids was done using an in-house spectral library with MS (and retention time), MS/MS information, and by searching the LIPID MAPS spectral database (
http://www.lipidmaps.org). MS/MS data were acquired in both negative and positive ion modes in order to maximize identification coverage. Additionally, some lipids were verified by injection of commercial standards. The identification was carried out in pooled cell extracts.

The peak area obtained for each lipid was normalized with lipid-class specific internal standards and with total content of protein. A (semi) quantitation was performed using lipid-class specific calibration curves. Pooled cell extracts were used for quality control, in addition to in-house plasma. The raw variation of the peak areas of internal standards in the samples was on average 15.3% and the RSD of retention times of identified lipids across all samples was on average 0.28%. The RSD of the concentrations of the identified lipids in QC samples and pooled extracts was on average 17.7%.

### Measurement of ceramides in Th17 and iTreg cells

#### Sample extraction

The frozen cell preps were defrosted on ice. The samples were extracted using a modified Folch method [71]. Briefly, 120 μL of cold (4 °C) extraction solvent (CHCl_3_: MeOH, (2:1 v/v) was added to the samples. The extraction solvent containing the following internal standards: C17 Lactosyl(β) ceramide (D18:1/17:0, 20 ppb), C17 Glucosyl(β) ceramide (D18:1/17:0, 20 ppb), C17 ceramide (D18:1/17:0, 20 ppb), C16 ceramide-d7 (d18:1-d7/16:0, 16,57 ppb), C18 ceramide-d7 (d18:1-d7/18:0, 8.75 ppb), C24 ceramide-d7 (d18:1-d7/24:0, 20 ppb), and C24:1 ceramide-d7 (d18:1-d7/24:1(15Z), 9,96 ppb). The samples were the vortexed briefly and left on ice for 30 minutes. The samples were then centrifuged (9400g, 5 min, 4 °C) and then 60 μL of the bottom layer was transfer to a clean mass spectrometry vial (2 mL). The samples were then stored at –80 °C.

#### Mass Spectrometry

The ceramides were quantified using a targeted multiple reaction monitoring (MRM) method using UHPLC as a separation technique. The LC separation was based on the global lipidomics method previously described [71]. Briefly, the UHPLC was a Exion AD (Sciex) integrated system. The samples were held in a cool box at 15 °C prior to the analysis. The needle was washed with both a 10% DCM in MeOH and ACN: MeOH: IPA: H_2_O (1:1:1:1 v/v/v/v) with 1% HCOOH for a total of 7.5 seconds each. The solvents were delivered using a quaternary solvent and a column oven (set to 50 °C). The separation was performed on an ACQUITY UHPLC BEH C18 column (2.1 mm × 100 mm, particle size 1.7 μm, Waters, Milford, MA, USA). The flow rate was set at 0.4 ml/min throughout the run with an injection volume of 1 μL. The following solvents were used for the gradient elution: Solvent A was H_2_O with 1% NH_4_Ac (1M) and HCOOH (0.1%) added. Solvent B was a mixture of ACN: IPA (1:1 v/v) with 1% NH_4_Ac (1M) and HCOOH (0.1%) added. The gradient was programmed as follows: 0 to 2 min 35-80% B, 2 to 7 min 80-100 % B, 7 to 14 min 100% B. The column was equilibrated with a 7min period of 35 % B prior to the next run. The mass spectrometer was a Sciex 5500 QTRAP (Sciex) set in scheduled MRM mode. The details of the MRM transitions can be seen in (**Table S1**). All lipids were identified for their fatty acid composition by MS/MS to confirm their exact identification, there was also a linear relationship between the increasing number of carbons in the lipid chain and its corresponding retention time. Due to the isobaric nature of sugars we were unable to differentiate Glc and Glc head groups. All data were integrated using the quantitation tool in MultiQuant (3.0.3), all peaks were manually checked. Any analytes which were over the concentration of the standard curve were diluted (1:25) with the same extraction solvent minus the internal standards. The quantification was performed using class-based internal standards and in the case of those ceramide species without an authentic standard in the standard curve mix, we used the closest related structure. The standard curve mixture contained: Glucosyl (β) C12 ceramide, Lactosyl (β) C12 ceramide, C18 ceramide (D18:1/18:1), C18:1 dihydroceramide (d18:0/18:1(9Z)) and was run at the following levels (all in ppb): 100, 80, 60, 50, 40, 30, 20, 10 for the C12 standards and 10, 8, 6, 5, 4, 3, 2,1 for all C18 standards.

### Statistical analysis

The lipidomic dataset was log_2_ transformed. Principal component analysis (PCA) was performed using *‘prcomp’* function included in the *‘stats’* R package, no outliers found. Sparse Partial Least Squares Discriminant Analysis (sPLS-DA) [39] of the T-cell subsets was performed using the *‘splsda’* function coded in the *‘mixOmics v6.3.2’* R package. In addition, several PLS-DA models between Th0 *vs.* Thp, Th1 *vs.* Th0, Th2 *vs.* Th0, Th17 *vs.* Th0 and iTreg *vs.* Th0 cells were developed and Variable Importance in Projection (VIP) scores [40] were estimated. The PLS-DA models were cross-validated [72] by 7-fold cross-validation and model diagnostics were generated using *‘perf’* function.

The multivariate PLS-DA analysis was followed by a univariate statistic; a paired t-test using the *’t.test’* function was performed to identify significant differences in the lipid intensities between T-cell subsets and their paired control (Th0). All lipids that passed one or more criteria for variable selection, *i.e.*, with the sPLS-DA model with an area under the ROC curve (AUC) >= 0.85; RC (>± 0.05), VIP scores > 1 or paired t-test; p-value < 0.05) were listed as significant. Initial p-values were subjected to multiple hypothesis testing correction, *vis-à-vis* False Discovery Rate (FDR) adjustment using the *‘p-adjust’* function. The *‘Heatmap.2’*, *‘boxplot’*, *’beanplot’*, *‘gplot’*, and *‘ggplot2’* libraries/packages were used for data visualization.

### Standardization of gene expression data for metabolic genes identification, T-cell-specific GEMs reconstruction and RM analysis

Lineage-specific normalized gene expression profiles of the human CD4^+^ Thp, Th1 and Th2 [73], Th17 [57], iTreg cells [56] and their paired control (Th0) were obtained from the literature and / or Gene Expression Omnibus (GEO) (https://www.ncbi.nlm.nih.gov/geo/) [74] with the accession numbers (Thp, Th1 & Th2, *GEO: GSE71646*), (Th17, *GEO: GSE52260*) and (iTreg, *GEO: GSE90570*) respectively. A list of differentially-expressed genes (FDR < 0.05) for each T-cell subset *vs.* Th0 was retrieved. The expression datasets were used for the identification of MGs and contextualization of HTimmR to T-cell-specific GEMs. In order to identify MGs, genes expressed in Thp, Th1, Th2, Th17 and iTreg cells were searched in the existing human metabolic reconstructions, *i.e.*, HMR2 [30], Edinburgh Human Metabolic Network (ETHMN) [75], RECON2 [76], and databases, *i.e.*, the Kyoto Encyclopedia of Genes and Genomes (KEGG)[31] and the Encyclopedia of Human Genes and Metabolism (HumanCyc) [32]. We found that HMR2 had the highest coverage of the MGs from our datasets. However, 160 MGs were missing in HMR2 and other metabolic reconstructions. Metabolic reactions (MRs) encoded by these genes were identified from the literature. These MRs were used for the reconstruction of HTimmR.

### Genome-scale metabolic reconstruction and modeling of human CD4^+^ T-cell subsets

#### Contextualization of HTimmR to CD4^+^ T helper-specific GEMs

Functional GEMs of Thp, Th1, Th2, Th17 and iTreg cells were developed by combining the E-Flux *[77]* and INIT [28] algorithms applied to HTimmR, used as a template model. HTimmR was contextualized for each CD4^+^ T-cell subset using lineage-specific gene expression data [28]. Subsequently, a draft GEM for each CD4^+^ T-cell subset that include the active metabolic reactions and their associated components (*e.g.*, metabolic genes, enzymes, metabolites and their interactions) was generated. The draft models were subjected to gap-filling by assigning basic metabolic tasks exhibited by normal, activated and / or differentiated T-cells (e.g., secretion of L-lactate). A quality control / sanity check [27] was performed using COnstraint-Based Reconstruction and Analysis Toolbox (COBRA toolbox v3.0) [78]. All the models were able to carry out basic metabolic tasks [30, 79]. Any blocked reactions were rectified or removed before knockout and flux analysis was performed. Mixed integer linear programming (MILP) was performed using *’MOSEK 8’* solver (licensed for the academic user) integrated in the RAVEN 2.0 suite [80]. Linear programming (LP) and optimization was performed using *‘ILOG-IBM CPLEX (version 128)’* solver.

#### Reporter metabolite analysis

The lineage-specific, differentially-expressed MGs identified in this study from different human CD4^+^ T-cell subsets was employed for the RM predictions. RM analysis was performed using the *‘reporterMetabolites’* function of the RAVEN 2.0 suite [80].

Overrepresentation analysis (ORA) of the RMs in the metabolic subsystems / pathways of human CD4^+^ T-cell subset was evaluated by a global hypergeometric test. RMs that were significantly (p < 0.05) altered between the T-cell subsets *vs.* Th0 were subjected to ORA. All the metabolic subsystems / pathways with (False Discovery Rate, FDR < 0.05) were listed.

#### Pathway knockout and essentiality analysis

An *in-silico* knockout (KO) analysis of sphingolipid pathways in human CD4^+^ Th17 cells was performed. Here, we evaluated the ability of a sphingolipid pathway to produce Cers in a wild type (WT) and KO models. In a WT model (no KO), flux through 8 different sphingolipid pathways which are directly associated with Cer biosynthesis were maximized one-by-one (as the objective function), and flux incurred by these pathways were recorded. Consequently, these fluxes were converted to (%), which depicts the relative contribution of the sphingolipid pathways towards total Cer production.

Next, we developed several KO models by iteratively removing a particular reaction / pathway (one at a time) from sphingolipid metabolism, and simultaneously estimating the maximum flux incurred by these 8 different pathways towards total Cer production. Likewise, the (%) of flux contributed by these pathways towards total Cer production was estimated in a KO model. The reaction / pathway KO analysis was performed using *‘removeReactions’* function coded in the COBRA toolbox (v3.0) [78]. All simulations were performed in MATLAB 2017b (Mathworks, Inc., Natick, MA, USA).

## SUPPLEMENTAL INFORMATION

Supplemental Information includes fourteen figures (**Figures S1–14**) and one table (**Table S1**).

## DATA AVAILABILITY

The untargeted and targeted lipidomic datasets generated in this study were submitted to the NIH Common Fund’s National Metabolomics Data Repository (NMDR) website, the Metabolomics Workbench (https://www.metabolomicsworkbench.org). The GEMs for human CD4^+^ T-cell subsets were deposited in BioModels [81] (www.ebi.ac.uk/biomodels/), and assigned an identifier MODEL2101270002.

## ACKNOWLEDGMENTS

We thank Marjo Hakkarinen and Sarita Heinonen (Turku Bioscience Center, University of Turku, Finland) for excellent technical assistance. We would like to acknowledge the Turku Metabolomics Centre and Biocenter Finland for their contribution to metabolomic analysis. We thank Dr. Aidan McGlinchey (School of Medical Sciences, Örebro University, Sweden) for assistance with editing the manuscript.

## AUTHOR CONTRIBUTIONS

MO proposed the study. RL and MO supervised the study. PS, UU, RL and MO assisted with the formulation of study design. PS performed GSMM and analyzed the multi-omics datasets. TH and AD supervised the metabolomics experiments. TL, MA, AD and TH acquired the metabolomics data. EK, VH, SBA, TB, MK and OR prepared and provided CD4^+^ T-cell subsets for metabolomics experiments. SBA and TB performed the siRNA knockdown experiments. PS and MO wrote the manuscript. All authors critically reviewed and approved the final manuscript.

MO is the guarantor of this work and, as such, had full access to all of the data in the study and takes responsibility for the integrity of the data and the accuracy of the data analysis.

## SUPPLEMENTAL INFORMATION

**Figure S1.**
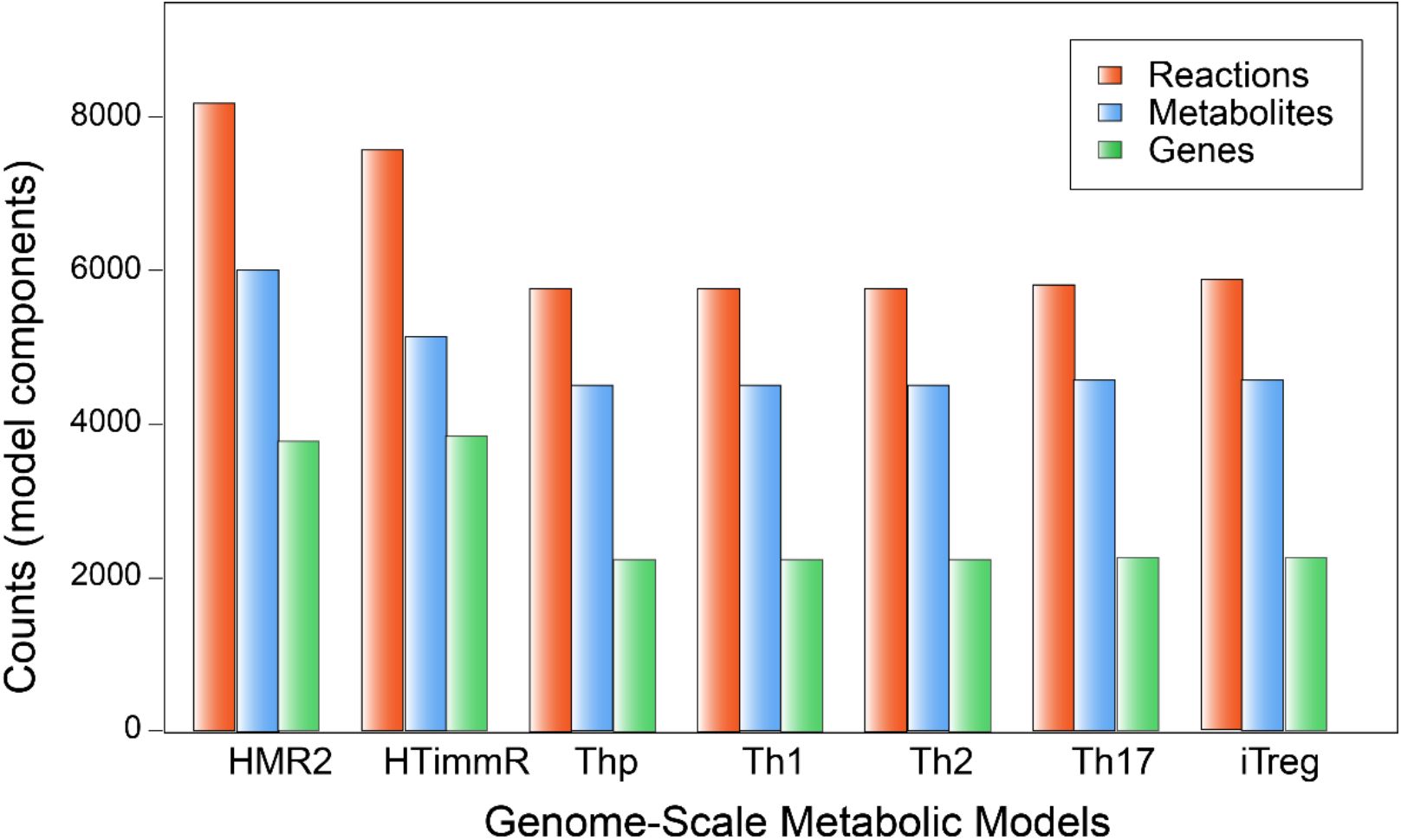
Genome-scale metabolic models (GEMs) of human CD4^+^ T-cell subsets. A bar plot comparing the number of reactions, metabolites and genes, included in the genome-scale metabolic models (GEMs) of the CD4^+^ T-cell subsets (Thp = T-naïve cells, T helpers: Th1, Th2, Th17, and iTreg cells). ‘HMR’ is Human Metabolic Reaction (GEM) [1] and ‘HTimmR’ is ‘Human T-immuno Reconstructor’ developed in this study.

**Figure S2.**
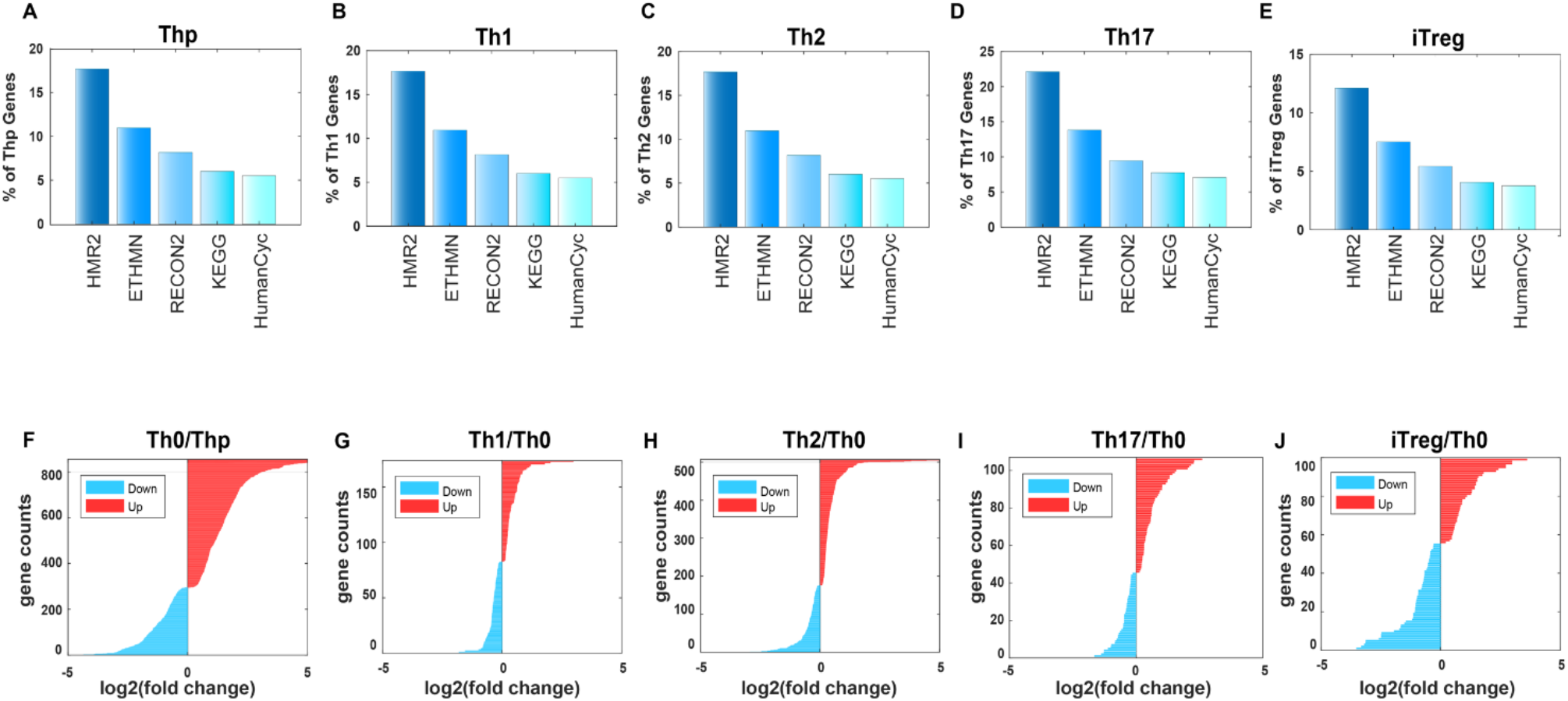
Metabolic genes and their differential expression in the human CD4^+^ T-cell subsets. A-E) It shows the percentage of genes in each CD4^+^ T-cell subset that are mapped to human metabolic reconstructions (HMR2, ETHMN, RECON) and pathway databases (KEGG and humanCyc). HMR2 has the highest coverage. ETHMN: Edinburg Human Metabolic Reconstruction [2], Recon2: a community-driven consensus metabolic reconstruction [3], HMR2: Human Metabolic Reaction, and databases [1], HumanCyc [4] and KEGG: Kyoto Encyclopedia of Genes and Genomes [5]. F-J) Log_2_ fold changes of differentially expressed (False discovery rate, FDR < 0.05) metabolic genes (MGs) in human CD4^+^ T-cell subsets (*vs.* Th0) during activation, and at 72 hours of polarization. Red and blue color denotes up- and downregulation of the MGs respectively.

**Figure S3.**
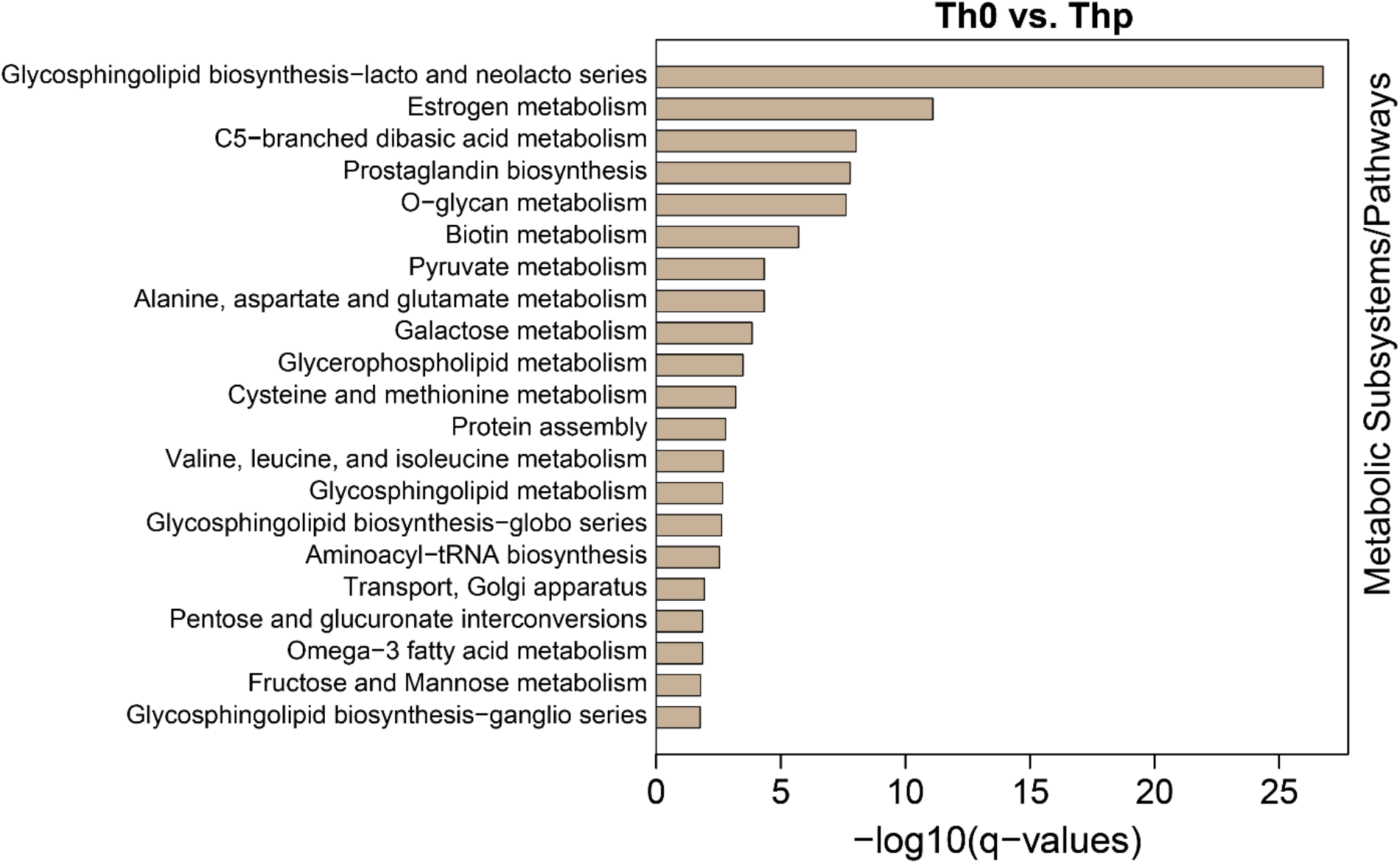
A bar plot showing overrepresented (hypergeometric test, q-values < 0.05) reporter pathways in the activated (Th0) *vs.* naïve (Thp) CD4^+^ T-cells.

**Figure S4.**
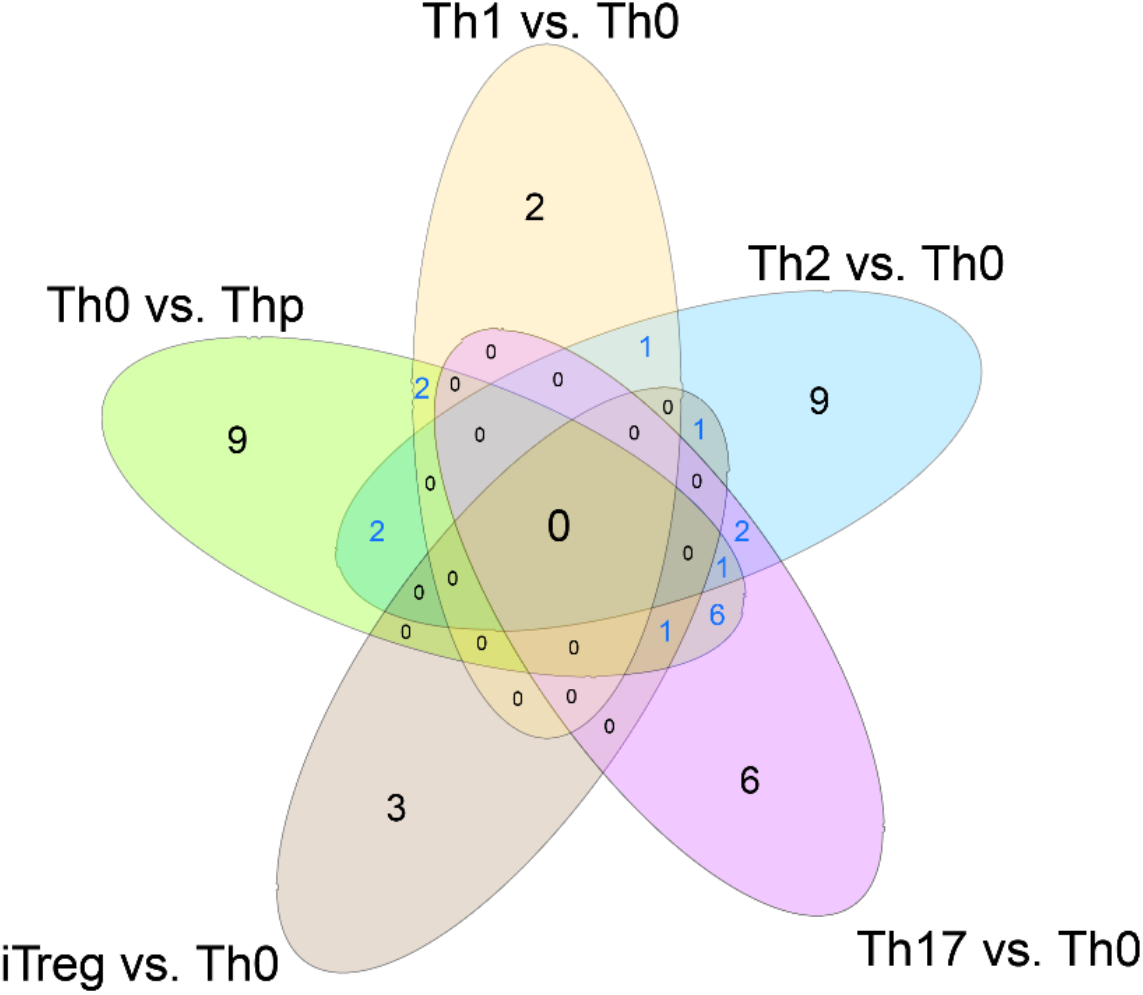
Venn diagram showing overrepresented metabolic subsystems / pathways which are common or unique among the CD4^+^ T-cell subsets when compared to Th0.

**Figure S5.**
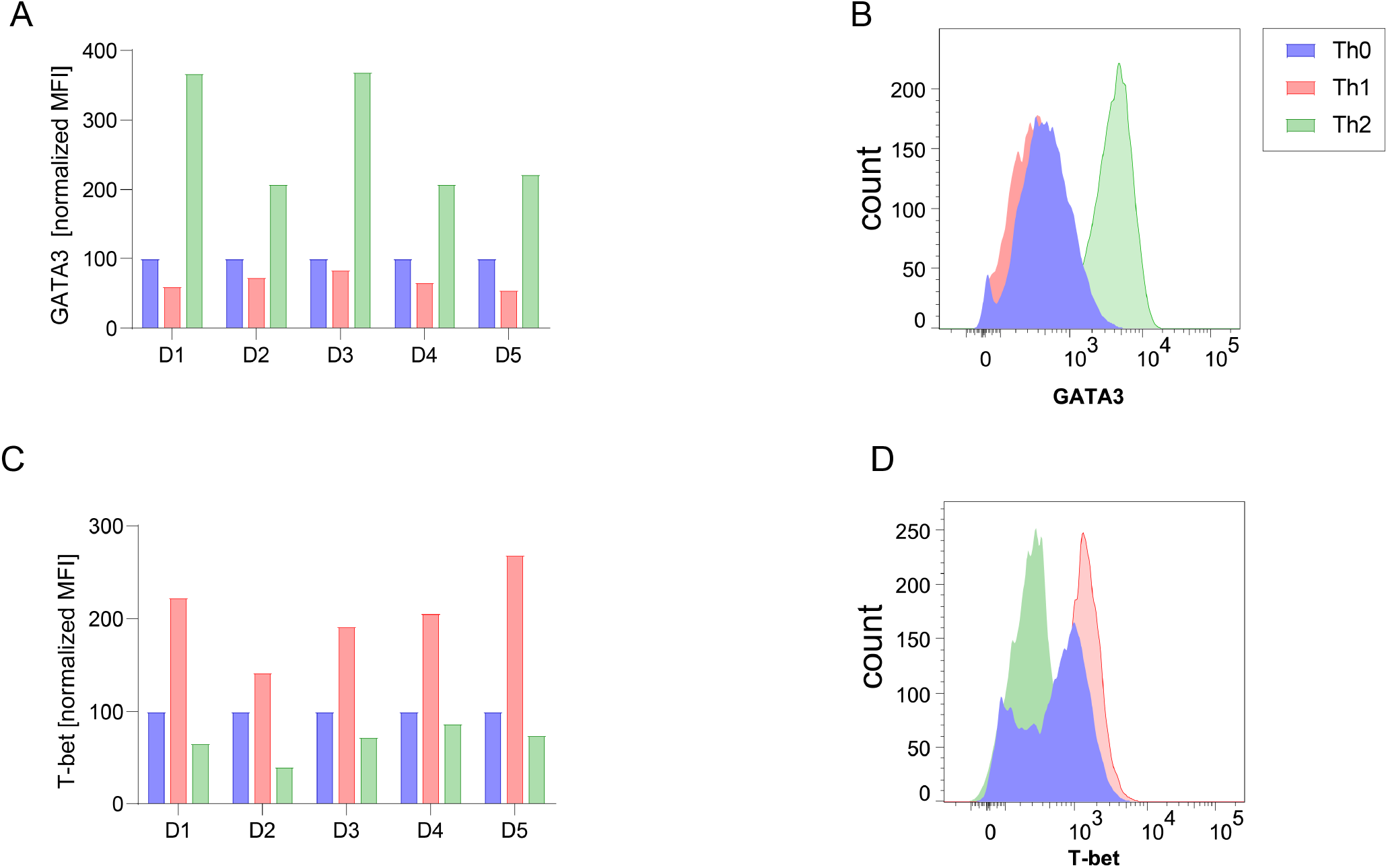
Intracellular protein expression of GATA3 and T-bet in 5 donors (D1-5) at 72 hours of Th0, Th1 and Th2 cell differentiation using flow cytometry.

**Figure S6.**
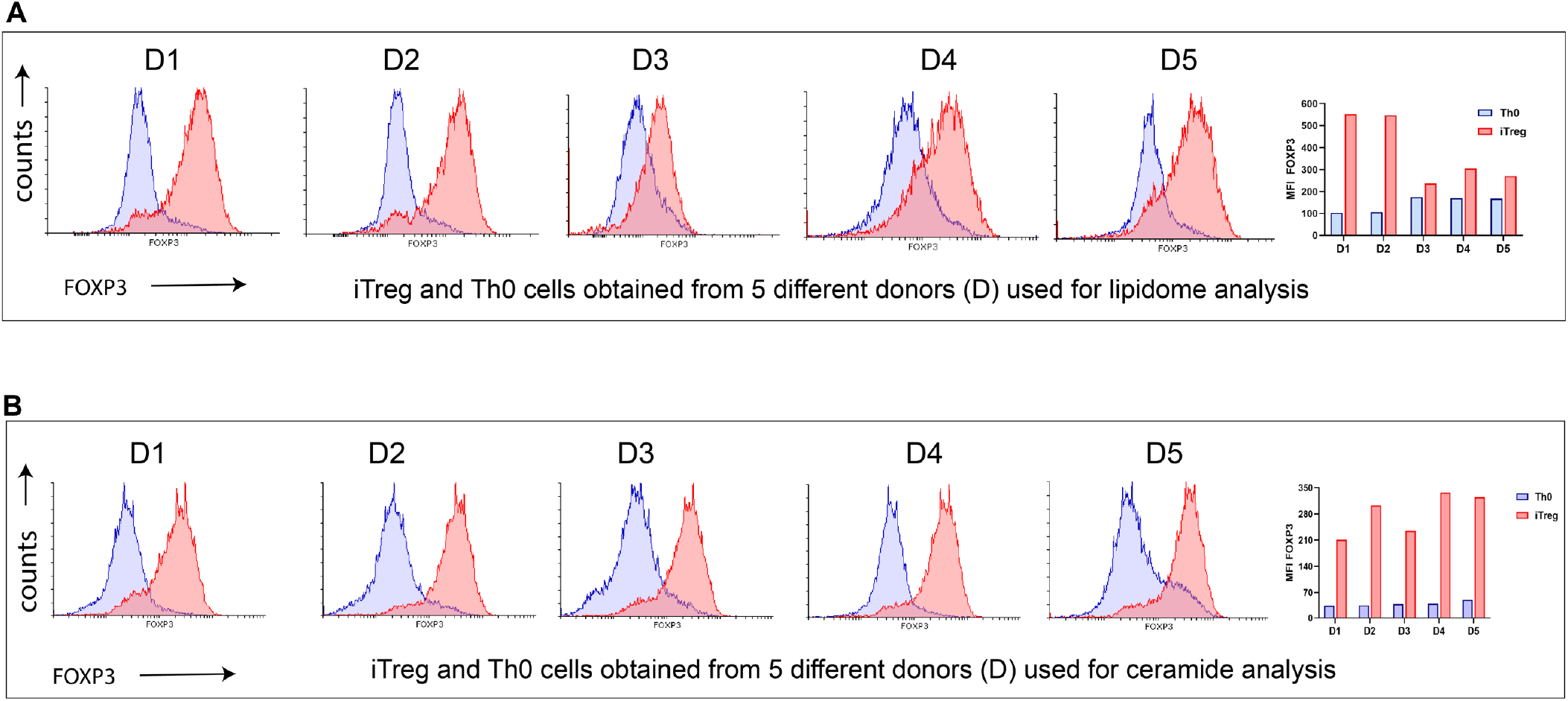
Intracellular FOXP3 expression in iTreg cells at 72 hours obtained from 5 donors (D1-5).

**Figure S7.**
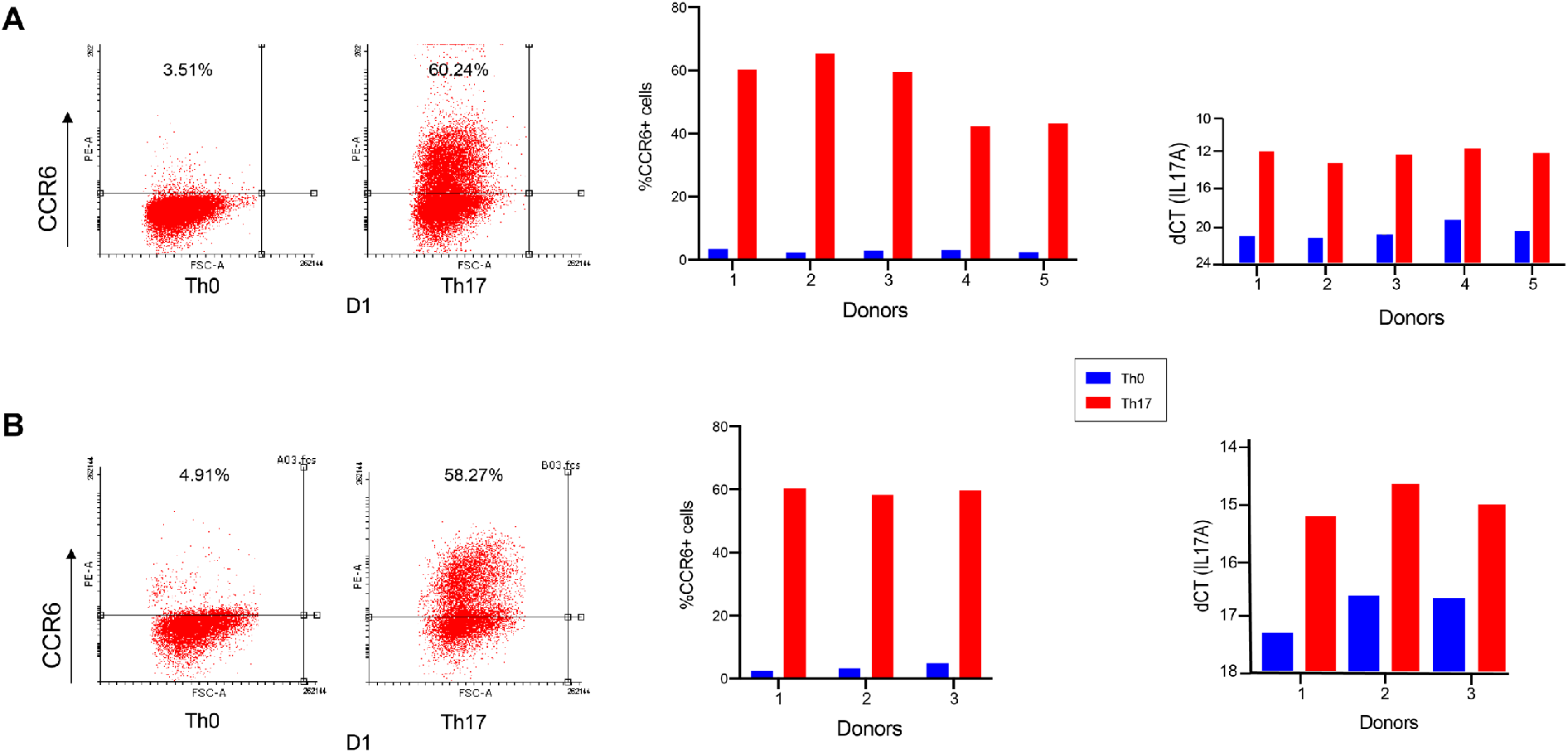
Flow cytometry analysis of CCR6 expression and TaqMan qPCR IL17A mRNA expression as the polarization readout for Th17 cells differentiated at 72h, and control Th0 cells. The data shown are representative dot plots, and bar plots for (A) five donor (D1-5) samples used for untargeted lipidomics and (B) three donor (D1-3) samples used for targeted ceramide measurements.

**Figure S8.**
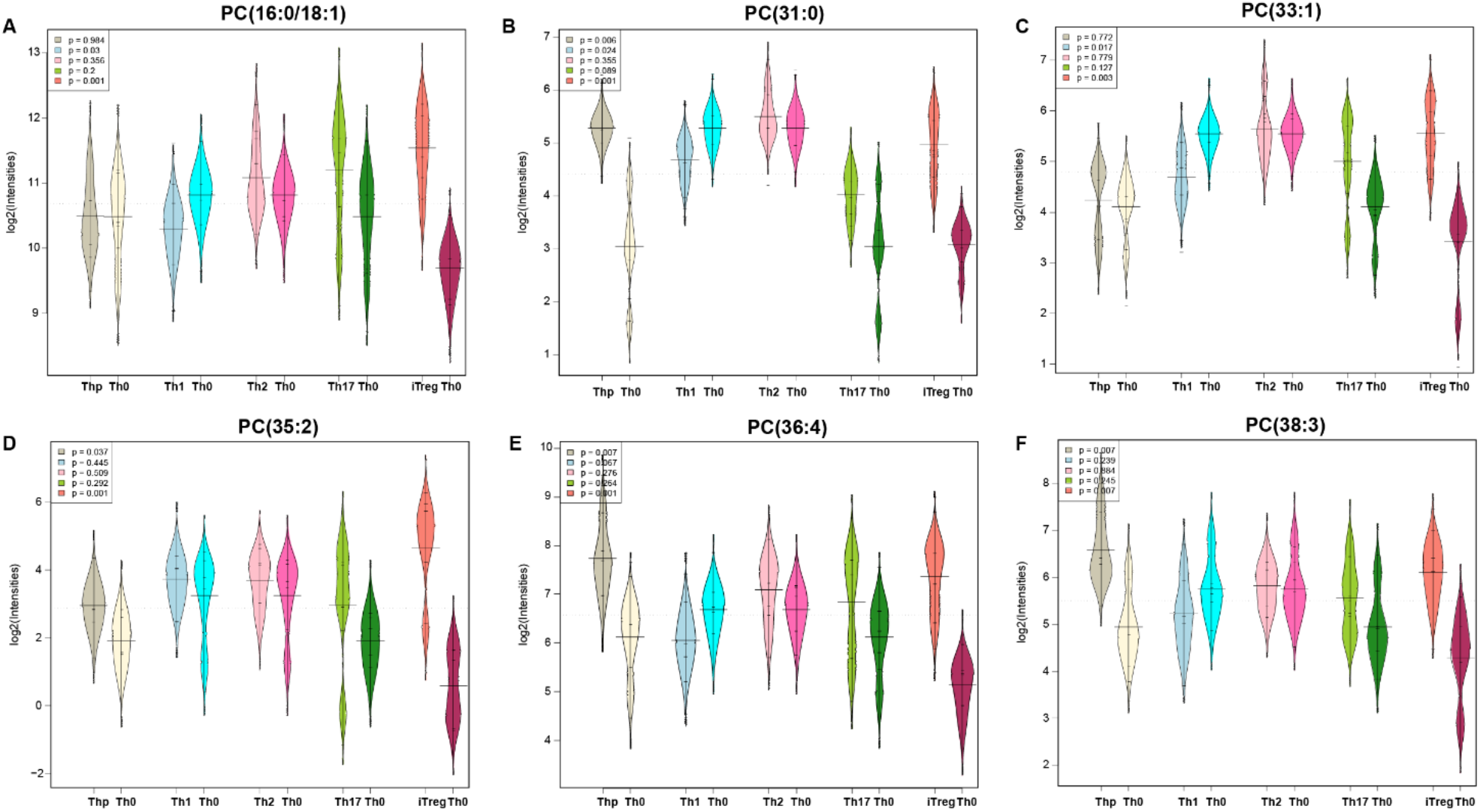
Beanplots showing the levels of PCs measured in T-naïve (Thp), Th1, Th2, Th17 and iTreg cells and their paired control (Th0) by non-targeted lipidomics, at 72 hours of polarization. >1 million cells were isolated from the umbilical cord of (n=5) healthy neonates. A significant difference is being determined by (paired t-test, p < 0.05). The black dashes in the bean plots represent the group mean. The legends show the p-values that are colored based on the groups/pairs.

**Figure S9.**
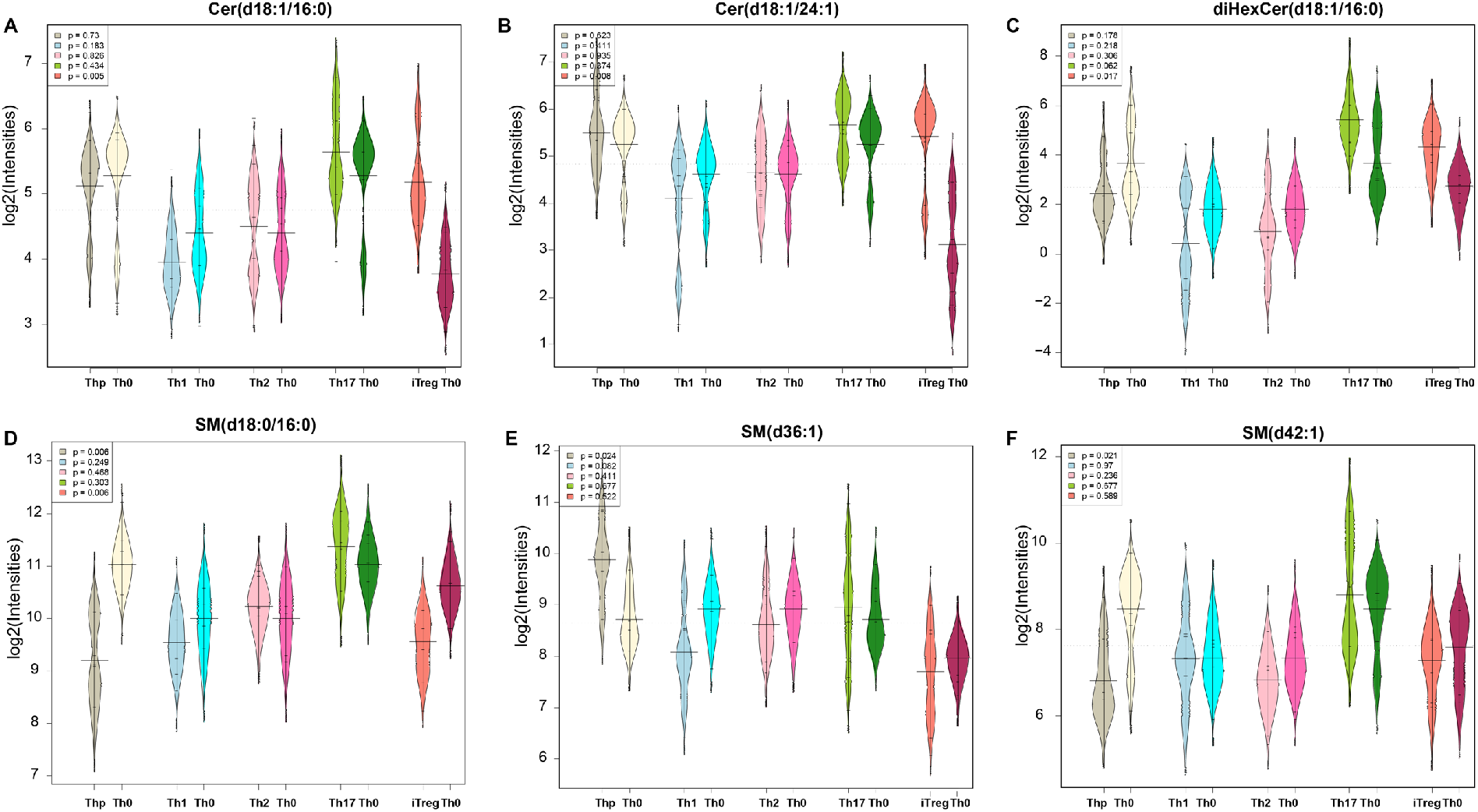
Beanplots showing the levels of Cers and SMs measured in Thp, Th1, Th2, Th17 and iTreg cells and their paired control (Th0) by non-targeted lipidomics, at 72 hours of polarization. A significant difference is being determined by (paired t-test, p < 0.05). The black dashes in the bean plots represent the group mean. The legends show the p-values that are colored based on the groups/pairs.

**Figure S10.**
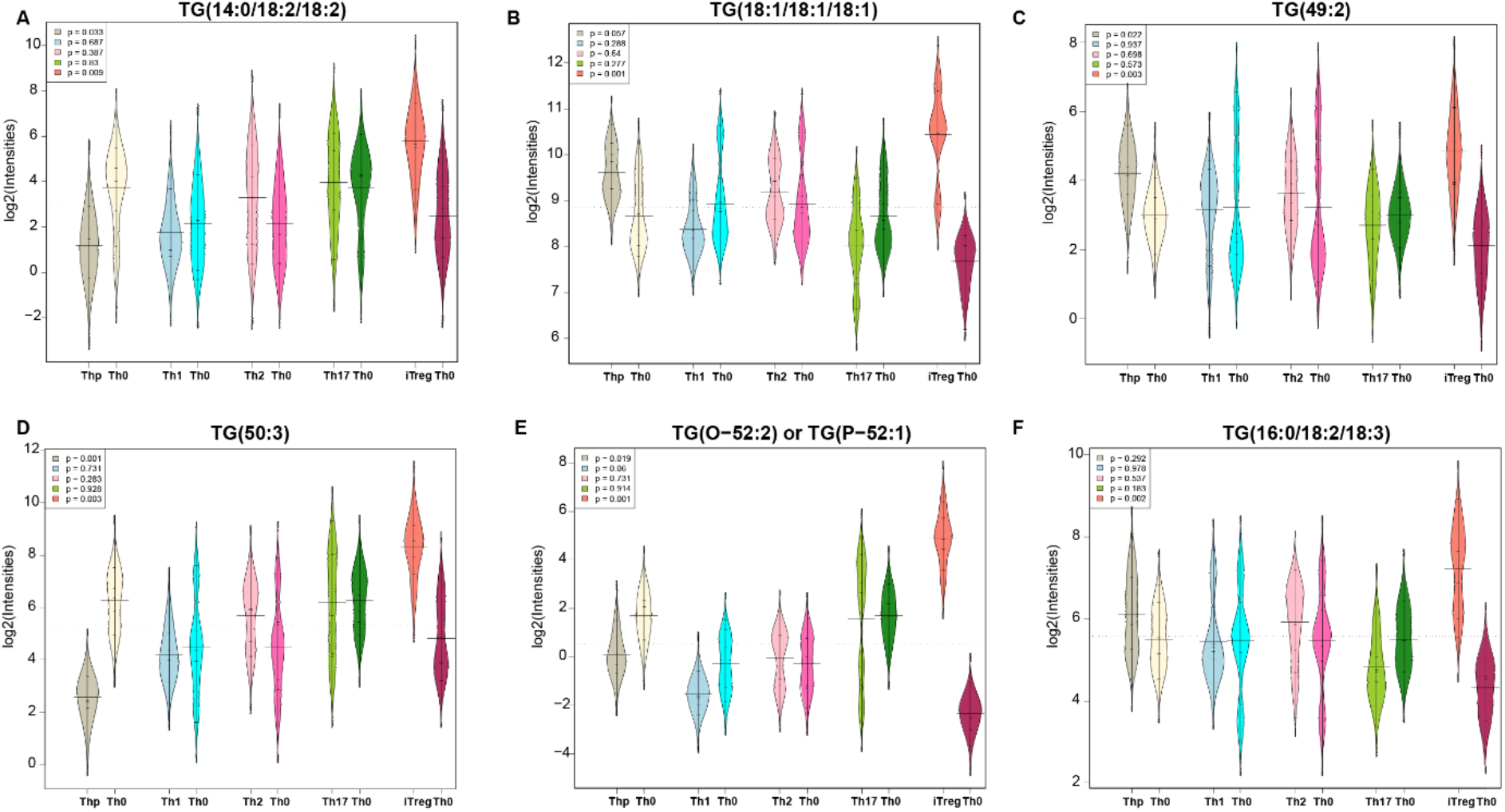
Beanplots showing the levels of TGs measured in Thp, Th1, Th2, Th17 and iTreg cells and their paired control (Th0) by non-targeted lipidomics, at 72 hours of polarization. A significant difference is being determined by (paired t-test, p < 0.05). The black dashes in the bean plots represent the group mean. The legends show the p-values that are colored based on the groups/pairs.

**Figure S11.**
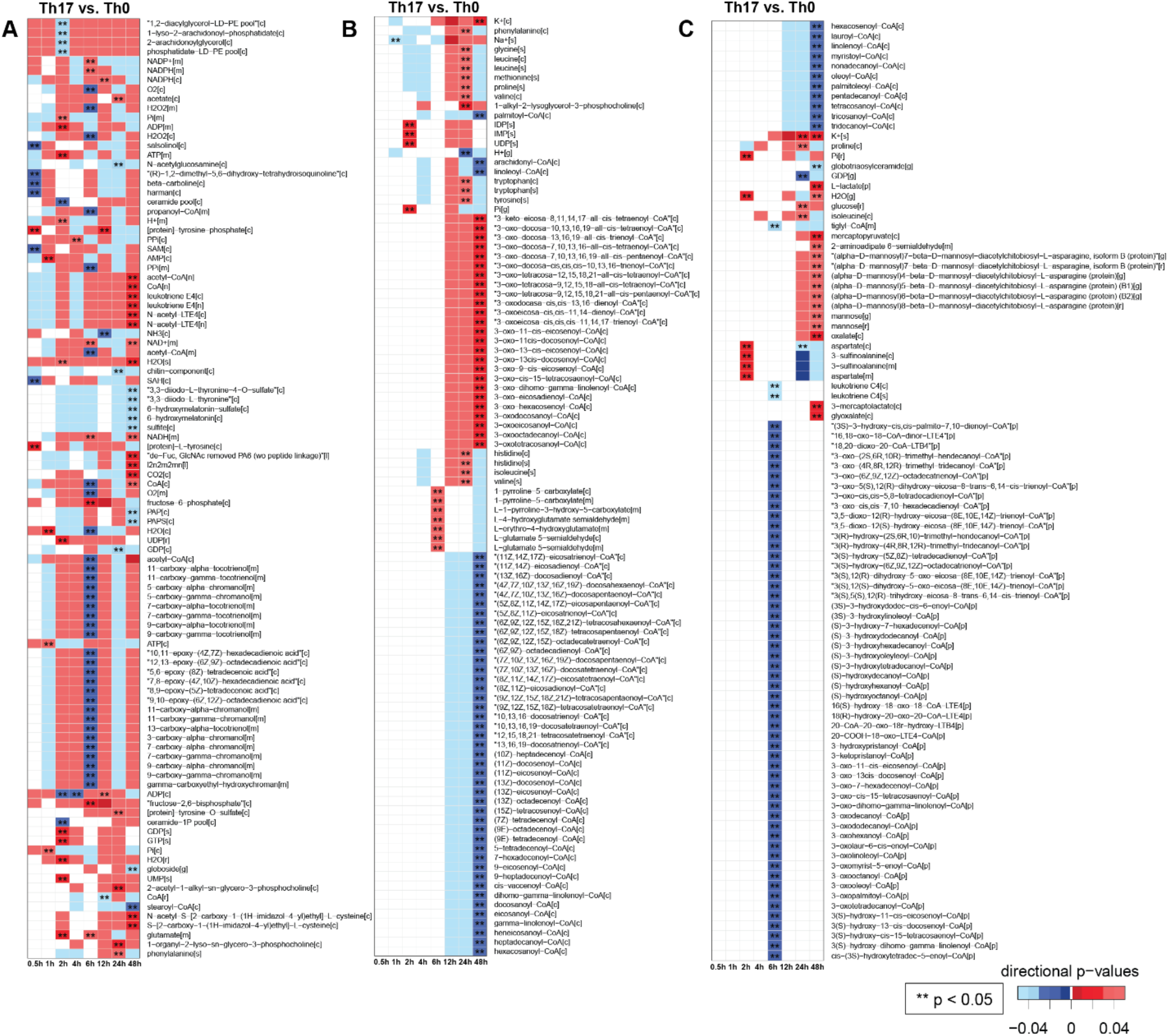
Reporter metabolites (RMs) of Th17 cells during the first 48 hours of differentiation. A-C) Heatmap of RMs (divided in multiple panels), that are significantly (p < 0.05) up- (red color) or downregulated (blue color) or remained unchanged or undetected (white color) in the Th17 cells as compared to their paired controls (Th0). The direction of regulation is given by the directional p-values [1, 6–8]. Stars (*) denotes various levels of significance *(**p < 0.05)*.

**Figure S12.**
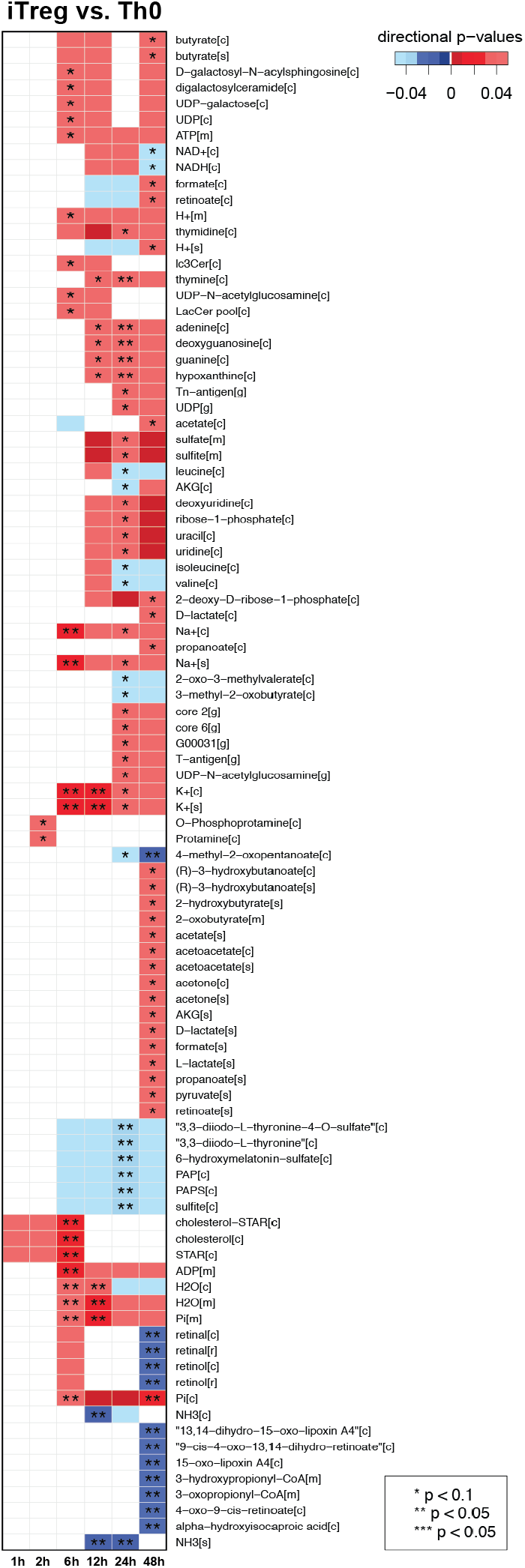
Reporter metabolites (RMs) of iTreg cells during the first 48 hours of differentiation. Heatmap of RMs that are significantly up- (red color) or downregulated (blue color) or remained unchanged or undetected (white color) in the iTreg cells as compared to their paired control (Th0). The direction of RM regulation is given by the directional p-values. Stars (*) denotes various levels of significance *(* p < 0.1, ** p < 0.05)*.

**Figure S13.**
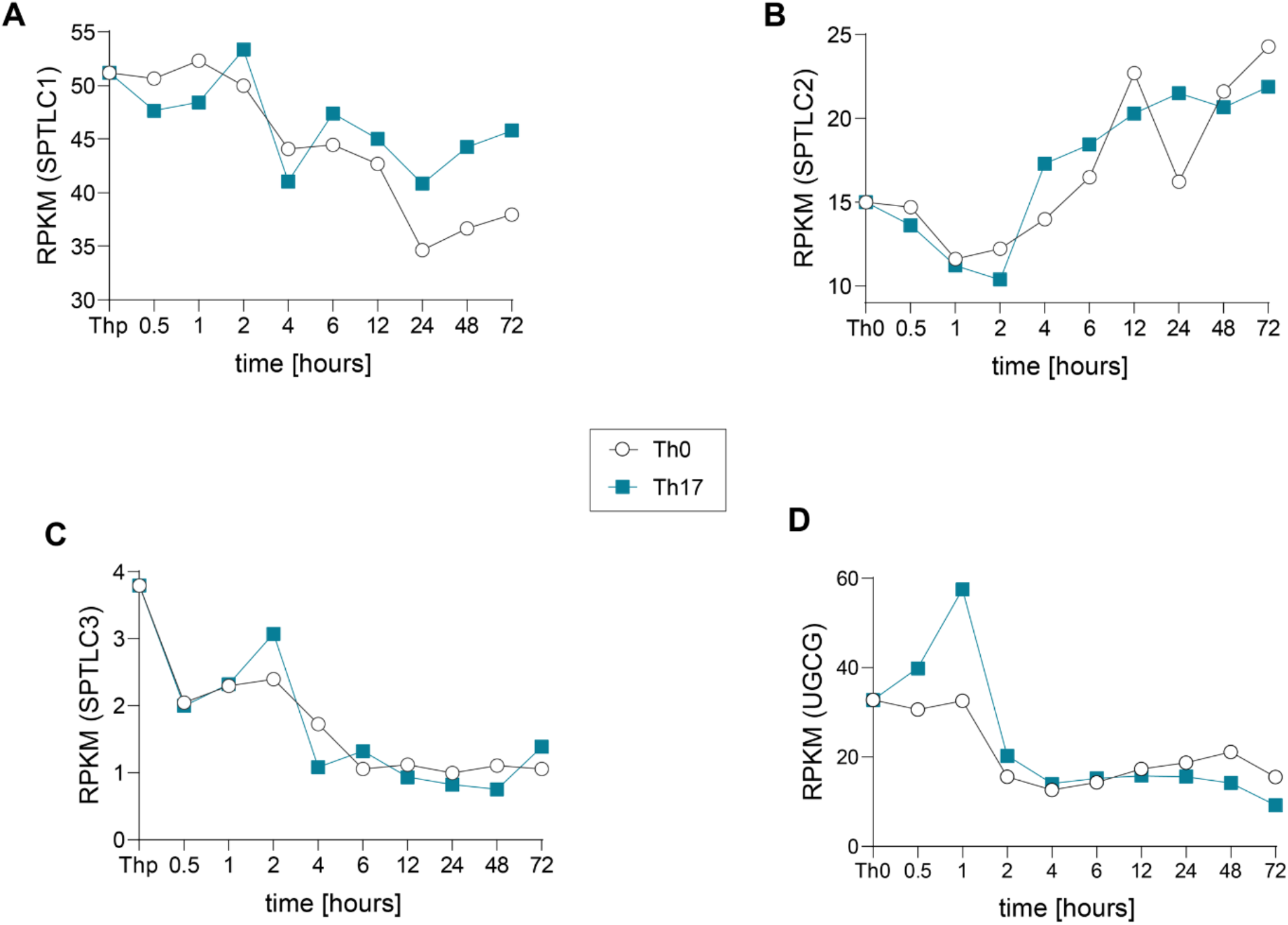
Expression profiles (RNA-Seq, mean ± SE) of *SPTLC1,2,3,* and *UGCG* genes in Th17 and Th0 cells obtained from cord blood of individuals across different time-points. ‘RPKM’ is Reads Per Kilobase Million.

**Figure S14.**
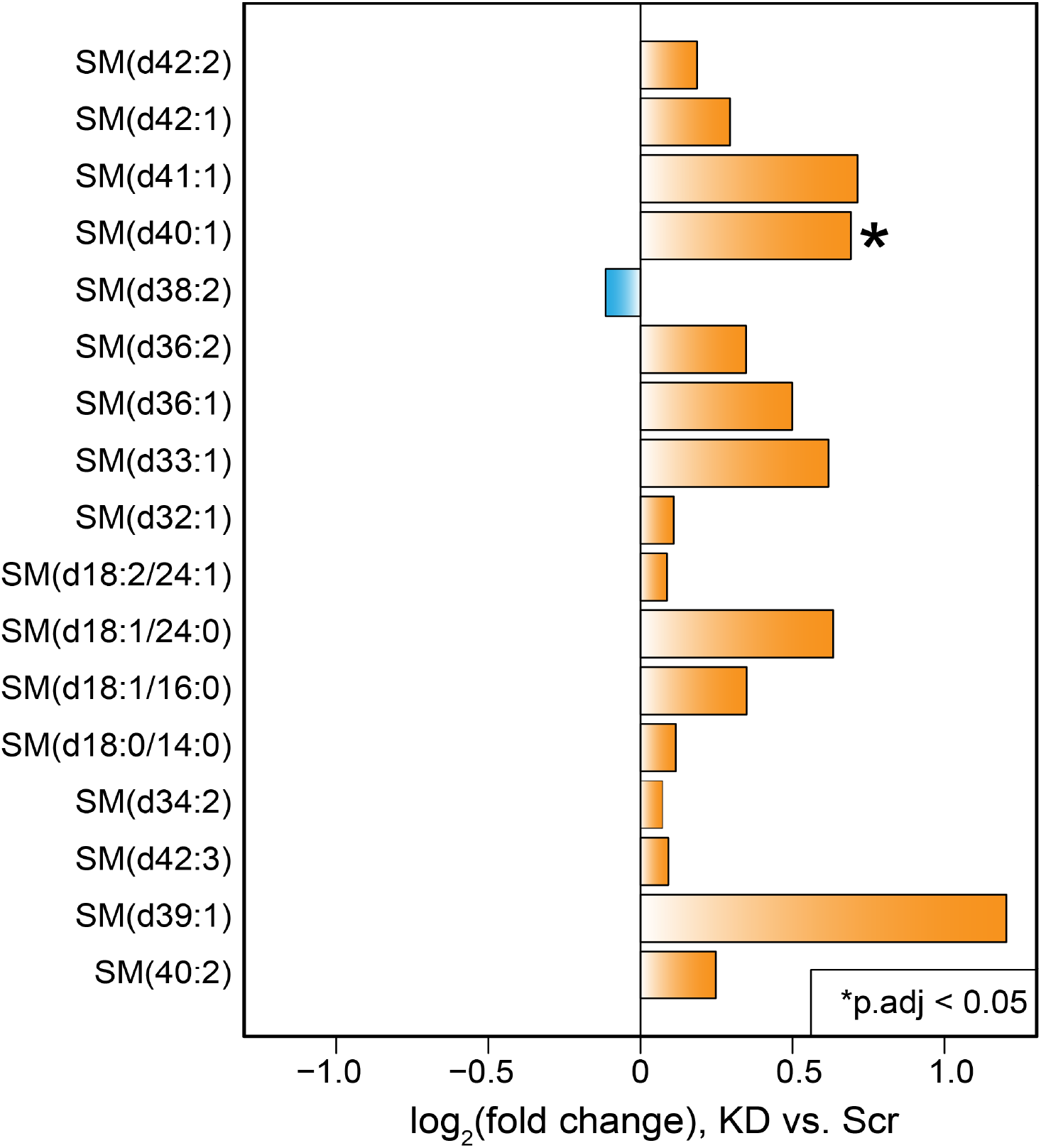
Targeted quantification of Sphingomyelins (SMs)in the *UGCG*-deficient Th17 cells. Bar plot showing log_2_ fold changes of the SMs measured in *UGCG*-deficient (KD) *vs.* control (Scr) Th17 cells at 72 hours (n=3).

**Table S1.** List of ceramides analyzed by targeted method.

| **Compound** | **Retention time (min)** | **Internal Standard** | **Precursor ion** | **Product ion** | **CE (+)** |
| --- | --- | --- | --- | --- | --- |
| **Cer(d18:1/12:0)** | 6.29 | Cer(d18:1-d7/16:0) | 482.4 | 264.2 | 45 |
| **Cer(d18:1/14:0)** | 6.86 | Cer(d18:1-d7/16:0) | 510.5 | 264.2 | 45 |
| **Cer(d18:1/16:0)** | 7.43 | Cer(d18:1-d7/16:0) | 538.5 | 264.2 | 45 |
| **Cer(d18:1/18:0)** | 7.93 | Cer(d18:1-d7/18:0) | 566.5 | 264.2 | 33 |
| **Cer(d18:1/20:0)** | 8.41 | Cer(d18:1-d7/18:0) | 594.6 | 264.2 | 33 |
| **Cer(d18:1/22:0)** | 8.9 | Cer(d18:1-d7/18:0) | 622.6 | 264.2 | 33 |
| **Cer(d18:1/24:0)** | 9.45 | Cer(d18:1-d7/24:0) | 650.6 | 264.2 | 36 |
| **Cer(d18:1/25:0)** | 9.69 | Cer(d18:1-d7/24:0) | 664.7 | 264.2 | 36 |
| **Cer(d18:1/26:0)** | 9.99 | Cer(d18:1-d7/24:0) | 678.7 | 264.2 | 36 |
| **Cer(d18:1/18:1(9Z))** | 7.43 | Cer(d18:1-d7/24:1(15Z)) | 564.5 | 264.2 | 29 |
| **Cer(d18:1/24:1)** | 8.79 | Cer(d18:1-d7/24:1(15Z)) | 658.6 | 264.201 | 38 |
| **Cer(d18:1/26:1)** | 9.37 | Cer(d18:1-d7/24:1(15Z)) | 676.6 | 264.2 | 38 |
| **HexCer(d18:1/12:0)** | 5.96 | HexCer(D18:1/17:0) | 644.5 | 264.2 | 45 |
| **HexCer(d18:1/16:0)** | 7.01 | HexCer(D18:1/17:0) | 700.5 | 264.2 | 45 |
| **HexCer(d18:1/18:0)** | 7.57 | HexCer(D18:1/17:0) | 728.1 | 264.2 | 43 |
| **HexCer(d18:1/20:0)** | 8.02 | HexCer(D18:1/17:0) | 756.6 | 264.2 | 45 |
| **HexCer(d18:1/22:0)** | 8.48 | HexCer(D18:1/17:0) | 784.7 | 264.2 | 45 |
| **HexCer(d18:1/23:0)** | 8.69 | HexCer(D18:1/17:0) | 798.7 | 264.2 | 45 |
| **HexCer(d18:1/24:0)** | 8.93 | HexCer(D18:1/17:0) | 812.7 | 264.2 | 45 |
| **HexCer(d18:1/18:1(9Z))** | 7.01 | HexCer(D18:1/17:0) | 726.6 | 264.2 | 45 |
| **HexCer(d18:1/24:1)** | 8.36 | HexCer(D18:1/17:0) | 810.7 | 264.2 | 45 |
| **diHexCer(d18:1/12:0)** | 5.79 | diHexCer(D18:1/17:0) | 806.5 | 264.2 | 45 |
| **diHexCer(d18:1/14:0)** | 6.24 | diHexCer(D18:1/17:0) | 834.6 | 264.2 | 53 |
| **diHexCer(d18:1/16:0)** | 6.83 | diHexCer(D18:1/17:0) | 862.6 | 264.2 | 53 |
| **diHexCer(d18:1/18:0)** | 7.4 | diHexCer(D18:1/17:0) | 890.2 | 264.2 | 52 |
| **diHexCer(d18:1/20:0)** | 10.52 | diHexCer(D18:1/17:0) | 918.7 | 264.2 | 53 |

## REFERENCES

1. Zhu J, Paul WE. CD4 T cells: fates, functions, and faults. Blood. 2008;112(5):1557–69. doi: 10.1182/blood-2008-05-078154.

2. Nurieva R, Wang J, Sahoo A. T-cell tolerance in cancer. Immunotherapy. 2013;5(5):513–31. doi:10.2217/imt.13.33.

3. Liblau RS, Wong FS, Mars LT, Santamaria P. Autoreactive CD8 T cells in organ-specific autoimmunity: emerging targets for therapeutic intervention. Immunity. 2002;17(1):1–6. doi:10.1016/s1074-7613(02)00338-2.

4. Macintyre AN, Rathmell JC. Activated lymphocytes as a metabolic model for carcinogenesis. Cancer & metabolism. 2013;1(1):5.

5. Chang C-H, Pearce EL. Emerging concepts in immunotherapy–T cell metabolism as a therapeutic target. Nature immunology. 2016;17(4):364.

6. Dimeloe S, Burgener AV, Grahlert J, Hess C. T-cell metabolism governing activation, proliferation and differentiation; a modular view. Immunology. 2017;150(1):35–44. doi:10.1111/imm.12655.

7. MacIver NJ, Michalek RD, Rathmell JC. Metabolic regulation of T lymphocytes. Annu Rev Immunol. 2013;31:259–83. doi:10.1146/annurev-immunol-032712-095956.

8. O’Shea JJ, Paul WE. Mechanisms underlying lineage commitment and plasticity of helper CD4+ T cells. Science. 2010;327(5969):1098–102. doi:10.1126/science.1178334.

9. Tuomela S, Lahesmaa R. Early T helper cell programming of gene expression in human. Semin Immunol. 2013;25(4):282–90. doi:10.1016/j.smim.2013.10.013.

10. Sugiura A, Rathmell JC. Metabolic Barriers to T Cell Function in Tumors. J Immunol. 2018;200(2):400–7. doi:10.4049/jimmunol.1701041.

11. Buck MD, O’Sullivan D, Pearce EL. T cell metabolism drives immunity. J Exp Med. 2015;212(9):1345–60. doi:10.1084/jem.20151159.

12. Pearce EL, Pearce EJ. Metabolic pathways in immune cell activation and quiescence. Immunity. 2013;38(4):633–43. doi:10.1016/j.immuni.2013.04.005.

13. Coloff JL, Mason EF, Altman BJ, Gerriets VA, Liu T, Nichols AN, et al. Akt requires glucose metabolism to suppress puma expression and prevent apoptosis of leukemic T cells. J Biol Chem. 2011;286(7):5921–33. doi:10.1074/jbc.M110.179101.

14. Calder PC. Fuel utilization by cells of the immune system. Proc Nutr Soc. 1995;54(1):65–82. doi:10.1079/pns19950038.

15. Barbi J, Pardoll D, Pan F. Metabolic control of the Treg/Th17 axis. Immunol Rev. 2013;252(1):52–77. doi:10.1111/imr.12029.

16. Powell JD, Delgoffe GM. The mammalian target of rapamycin: linking T cell differentiation, function, and metabolism. Immunity. 2010;33(3):301–11. doi:10.1016/j.immuni.2010.09.002.

17. Dang EV, Barbi J, Yang HY, Jinasena D, Yu H, Zheng Y, et al. Control of T(H)17/T(reg) balance by hypoxia-inducible factor 1. Cell. 2011;146(5):772–84. doi:10.1016/j.cell.2011.07.033.

18. Carr EL, Kelman A, Wu GS, Gopaul R, Senkevitch E, Aghvanyan A, et al. Glutamine uptake and metabolism are coordinately regulated by ERK/MAPK during T lymphocyte activation. J Immunol. 2010;185(2):1037–44. doi:10.4049/jimmunol.0903586.

19. Geiger R, Rieckmann JC, Wolf T, Basso C, Feng Y, Fuhrer T, et al. L-Arginine Modulates T Cell Metabolism and Enhances Survival and Anti-tumor Activity. Cell. 2016;167(3):829–42 e13. doi:10.1016/j.cell.2016.09.031.

20. Berod L, Friedrich C, Nandan A, Freitag J, Hagemann S, Harmrolfs K, et al. De novo fatty acid synthesis controls the fate between regulatory T and T helper 17 cells. Nature medicine. 2014;20(11):1327–33.

21. Pearce EL, Poffenberger MC, Chang C-H, Jones RG. Fueling immunity: insights into metabolism and lymphocyte function. Science. 2013;342(6155):1242454.

22. Gerriets VA, Rathmell JC. Metabolic pathways in T cell fate and function. Trends Immunol. 2012;33(4):168–73. doi:10.1016/j.it.2012.01.010.

23. Chang CH, Curtis JD, Maggi LB, Jr., Faubert B, Villarino AV, O’Sullivan D, et al. Posttranscriptional control of T cell effector function by aerobic glycolysis. Cell. 2013;153(6):1239–51. doi:10.1016/j.cell.2013.05.016.

24. O’Sullivan D, Pearce EL. Targeting T cell metabolism for therapy. Trends in immunology. 2015;36(2):71–80.

25. Orth JD, Thiele I, Palsson BØ. What is flux balance analysis? Nature biotechnology. 2010;28(3):245–8.

26. Sen P, Kemppainen E, Orešič M. Perspectives on Systems Modelling of Human Peripheral Blood Mononuclear Cells. Frontiers in molecular biosciences. 2017;4:96.

27. Thiele I, Palsson BO. A protocol for generating a high-quality genome-scale metabolic reconstruction. Nat Protoc. 2010;5(1):93–121. doi:10.1038/nprot.2009.203.

28. Agren R, Bordel S, Mardinoglu A, Pornputtapong N, Nookaew I, Nielsen J. Reconstruction of genome-scale active metabolic networks for 69 human cell types and 16 cancer types using INIT. PLoS Comput Biol. 2012;8(5):e1002518.

29. Sen P, Dickens AM, Lopez-Bascon MA, Lindeman T, Kemppainen E, Lamichhane S, et al. Metabolic alterations in immune cells associate with progression to type 1 diabetes. Diabetologia. 2020;63(5):1017–31. doi:10.1007/s00125-020-05107-6.

30. Mardinoglu A, Agren R, Kampf C, Asplund A, Uhlen M, Nielsen J. Genome-scale metabolic modelling of hepatocytes reveals serine deficiency in patients with non-alcoholic fatty liver disease. Nature communications. 2014;5.

31. Kanehisa M, Goto S. KEGG: kyoto encyclopedia of genes and genomes. Nucleic acids research. 2000;28(1):27–30.

32. Trupp M, Altman T, Fulcher CA, Caspi R, Krummenacker M, Paley S, et al. Beyond the genome (BTG) is a (PGDB) pathway genome database: HumanCyc. Genome biology. 2010;11(1):O12.

33. Cakir T, Patil KR, Onsan Z, Ulgen KO, Kirdar B, Nielsen J. Integration of metabolome data with metabolic networks reveals reporter reactions. Mol Syst Biol. 2006;2:50. doi:10.1038/msb4100085.

34. Patil KR, Nielsen J. Uncovering transcriptional regulation of metabolism by using metabolic network topology. Proceedings of the National Academy of Sciences of the United States of America. 2005;102(8):2685–9. doi:10.1073/pnas.0406811102.

35. Ryan DG, O’Neill LA. Krebs cycle rewired for macrophage and dendritic cell effector functions. FEBS letters. 2017;591(19):2992–3006.

36. Hooftman A, O’Neill LAJ. The Immunomodulatory Potential of the Metabolite Itaconate. Trends Immunol. 2019;40(8):687–98. doi:10.1016/j.it.2019.05.007.

37. Dantzer R. Role of the Kynurenine Metabolism Pathway in Inflammation-Induced Depression: Preclinical Approaches. Curr Top Behav Neurosci. 2017;31:117–38. doi:10.1007/7854_2016_6.

38. Poffenberger MC, Jones RG. Amino acids fuel T cell-mediated inflammation. Immunity. 2014;40(5):635–7.

39. Le Cao KA, Boitard S, Besse P. Sparse PLS discriminant analysis: biologically relevant feature selection and graphical displays for multiclass problems. BMC Bioinformatics. 2011;12:253. doi:10.1186/1471-2105-12-253.

40. Farrés M, Platikanov S, Tsakovski S, Tauler R. Comparison of the variable importance in projection (VIP) and of the selectivity ratio (SR) methods for variable selection and interpretation. J Chemom. 2015;29(10):528–36.

41. Zhang T, de Waard AA, Wuhrer M, Spaapen RM. The Role of Glycosphingolipids in Immune Cell Functions. Front Immunol. 2019;10:90. doi:10.3389/fimmu.2019.00090.

42. Apostolidis SA, Rodríguez-Rodríguez N, Suárez-Fueyo A, Dioufa N, Ozcan E, Crispín JC, et al. Phosphatase PP2A is requisite for the function of regulatory T cells. Nature immunology. 2016;17(5):556.

43. Hanada K. Serine palmitoyltransferase, a key enzyme of sphingolipid metabolism. Biochim Biophys Acta. 2003;1632(1-3):16–30. doi:10.1016/s1388-1981(03)00059-3.

44. Hornemann T, Penno A, Rutti MF, Ernst D, Kivrak-Pfiffner F, Rohrer L, et al. The SPTLC3 subunit of serine palmitoyltransferase generates short chain sphingoid bases. J Biol Chem. 2009;284(39):26322–30. doi:10.1074/jbc.M109.023192.

45. Alam S, Fedier A, Kohler RS, Jacob F. Glucosylceramide synthase inhibitors differentially affect expression of glycosphingolipids. Glycobiology. 2015;25(4):351–6. doi:10.1093/glycob/cwu187.

46. Newton R, Priyadharshini B, Turka LA. Immunometabolism of regulatory T cells. Nat Immunol. 2016;17(6):618–25. doi:10.1038/ni.3466.

47. Klysz D, Tai X, Robert PA, Craveiro M, Cretenet G, Oburoglu L, et al. Glutamine-dependent α-ketoglutarate production regulates the balance between T helper 1 cell and regulatory T cell generation. Sci Signal. 2015;8(396):ra97–ra.

48. Gault CR, Obeid LM, Hannun YA. An overview of sphingolipid metabolism: from synthesis to breakdown. Adv Exp Med Biol. 2010;688:1–23. doi:10.1007/978-1-4419-6741-1_1.

49. Adam D, Heinrich M, Kabelitz D, Schütze S. Ceramide: does it matter for T cells? Trends in immunology. 2002;23(1):1–4.

50. Arvey A, van der Veeken J, Samstein RM, Feng Y, Stamatoyannopoulos JA, Rudensky AY. Inflammation-induced repression of chromatin bound by the transcription factor Foxp3 in regulatory T cells. Nat Immunol. 2014;15(6):580–7. doi:10.1038/ni.2868.

51. Apostolidis SA, Rauen T, Hedrich CM, Tsokos GC, Crispín JC. Protein phosphatase 2A enables expression of interleukin 17 (IL-17) through chromatin remodeling. Journal of Biological Chemistry. 2013;288(37):26775–84.

52. Xu Q, Jin X, Zheng M, Rohila D, Fu G, Wen Z, et al. Phosphatase PP2A is essential for TH17 differentiation. Proceedings of the National Academy of Sciences. 2019;116(3):982–7.

53. Bai A, Moss A, Kokkotou E, Usheva A, Sun X, Cheifetz A, et al. CD39 and CD161 modulate Th17 responses in Crohn’s disease. J Immunol. 2014;193(7):3366–77. doi:10.4049/jimmunol.1400346.

54. Bai A, Robson S. Beyond ecto-nucleotidase: CD39 defines human Th17 cells with CD161. Purinergic Signal. 2015;11(3):317–9. doi:10.1007/s11302-015-9457-4.

55. Bai A, Guo Y. Acid sphingomyelinase mediates human CD4(+) T-cell signaling: potential roles in T-cell responses and diseases. Cell Death Dis. 2017;8(7):e2963. doi:10.1038/cddis.2017.360.

56. Ubaid U, Andrabi SBA, Tripathi SK, Dirasantha O, Kanduri K, Rautio S, et al. Transcriptional Repressor HIC1 Contributes to Suppressive Function of Human Induced Regulatory T Cells. Cell Rep. 2018;22(8):2094–106. doi:10.1016/j.celrep.2018.01.070.

57. Tuomela S, Rautio S, Ahlfors H, Oling V, Salo V, Ullah U, et al. Comparative analysis of human and mouse transcriptomes of Th17 cell priming. Oncotarget. 2016;7(12):13416–28. doi:10.18632/oncotarget.7963.

58. Sattar N, McInnes IB, McMurray JJV. Obesity Is a Risk Factor for Severe COVID-19 Infection: Multiple Potential Mechanisms. Circulation. 2020;142(1):4–6. doi:10.1161/CIRCULATIONAHA.120.047659.

59. Virtue S, Vidal-Puig A. Adipose tissue expandability, lipotoxicity and the Metabolic Syndrome--an allostatic perspective. Biochim Biophys Acta. 2010;1801(3):338–49. doi:10.1016/j.bbalip.2009.12.006.

60. Wang F, Hou H, Luo Y, Tang G, Wu S, Huang M, et al. The laboratory tests and host immunity of COVID-19 patients with different severity of illness. JCI Insight. 2020;5(10). doi:10.1172/jci.insight.137799.

61. Wu D, Yang XO. TH17 responses in cytokine storm of COVID-19: An emerging target of JAK2 inhibitor Fedratinib. J Microbiol Immunol Infect. 2020;53(3):368–70. doi:10.1016/j.jmii.2020.03.005.

62. Khan MM, Ullah U, Khan MH, Kong L, Moulder R, Valikangas T, et al. CIP2A Constrains Th17 Differentiation by Modulating STAT3 Signaling. iScience. 2020;23(3):100947. doi:10.1016/j.isci.2020.100947.

63. Tripathi SK, Chen Z, Larjo A, Kanduri K, Nousiainen K, Aijo T, et al. Genome-wide Analysis of STAT3-Mediated Transcription during Early Human Th17 Cell Differentiation. Cell Rep. 2017;19(9):1888–901. doi:10.1016/j.celrep.2017.05.013.

64. Hawkins RD, Larjo A, Tripathi SK, Wagner U, Luu Y, Lonnberg T, et al. Global chromatin state analysis reveals lineage-specific enhancers during the initiation of human T helper 1 and T helper 2 cell polarization. Immunity. 2013;38(6):1271–84. doi:10.1016/j.immuni.2013.05.011.

65. Pedersen HK, Forslund SK, Gudmundsdottir V, Petersen AØ, Hildebrand F, Hyötyläinen T, et al. A computational framework to integrate high-throughput ‘-omics’ datasets for the identification of potential mechanistic links. Nature protocols. 2018:1.

66. Bradford MM. A rapid and sensitive method for the quantitation of microgram quantities of protein utilizing the principle of protein-dye binding. Anal Biochem. 1976;72:248–54.

67. Nygren H, Seppanen-Laakso T, Castillo S, Hyotylainen T, Oresic M. Liquid chromatography-mass spectrometry (LC-MS)-based lipidomics for studies of body fluids and tissues. Methods Mol Biol. 2011;708:247–57. doi:10.1007/978-1-61737-985-7_15.

68. Pedersen HK, Forslund SK, Gudmundsdottir V, Petersen AO, Hildebrand F, Hyotylainen T, et al. A computational framework to integrate high-throughput ‘-omics’ datasets for the identification of potential mechanistic links. Nat Protoc. 2018;13(12):2781–800. doi:10.1038/s41596-018-0064-z.

69. Pluskal T, Castillo S, Villar-Briones A, Oresic M. MZmine 2: modular framework for processing, visualizing, and analyzing mass spectrometry-based molecular profile data. BMC Bioinform. 2010;11:395. doi:10.1186/1471-2105-11-395.

70. Pluskal T, Castillo S, Villar-Briones A, Orešič M. MZmine 2: modular framework for processing, visualizing, and analyzing mass spectrometry-based molecular profile data. BMC bioinformatics. 2010;11(1):1.

71. Sen P, Carlsson C, Virtanen SM, Simell S, Hyoty H, Ilonen J, et al. Persistent Alterations in Plasma Lipid Profiles Before Introduction of Gluten in the Diet Associated With Progression to Celiac Disease. Clin Transl Gastroenterol. 2019;10(5):1–10. doi:10.14309/ctg.0000000000000044.

72. Westerhuis JA, Hoefsloot HC, Smit S, Vis DJ, Smilde AK, van Velzen EJ, et al. Assessment of PLSDA cross validation. Metabolomics. 2008;4(1):81–9.

73. Kanduri K, Tripathi S, Larjo A, Mannerstrom H, Ullah U, Lund R, et al. Identification of global regulators of T-helper cell lineage specification. Genome Med. 2015;7:122. doi:10.1186/s13073-015-0237-0.

74. Edgar R, Domrachev M, Lash AE. Gene Expression Omnibus: NCBI gene expression and hybridization array data repository. Nucleic Acids Res. 2002;30(1):207–10. doi:10.1093/nar/30.1.207.

75. Ma H, Sorokin A, Mazein A, Selkov A, Selkov E, Demin O, et al. The Edinburgh human metabolic network reconstruction and its functional analysis. Mol Syst Biol. 2007;3:135. doi:10.1038/msb4100177.

76. Thiele I, Swainston N, Fleming RM, Hoppe A, Sahoo S, Aurich MK, et al. A community-driven global reconstruction of human metabolism. Nat Biotechnol. 2013;31(5):419–25. doi:10.1038/nbt.2488.

77. Colijn C, Brandes A, Zucker J, Lun DS, Weiner B, Farhat MR, et al. Interpreting expression data with metabolic flux models: predicting Mycobacterium tuberculosis mycolic acid production. PLoS computational biology. 2009;5(8):e1000489.

78. Heirendt L, Arreckx S, Pfau T, Mendoza SN, Richelle A, Heinken A, et al. Creation and analysis of biochemical constraint-based models: the COBRA Toolbox v3. 0. arXiv preprint arXiv:171004038. 2017.

79. Mardinoglu A, Agren R, Kampf C, Asplund A, Nookaew I, Jacobson P, et al. Integration of clinical data with a genome-scale metabolic model of the human adipocyte. Mol Syst Biol. 2013;9:649. doi:10.1038/msb.2013.5.

80. Wang H, Marcisauskas S, Sanchez BJ, Domenzain I, Hermansson D, Agren R, et al. RAVEN 2.0: A versatile toolbox for metabolic network reconstruction and a case study on Streptomyces coelicolor. PLoS Comput Biol. 2018;14(10):e1006541. doi:10.1371/journal.pcbi.1006541.

81. Malik-Sheriff RS, Glont M, Nguyen TVN, Tiwari K, Roberts MG, Xavier A, et al. BioModels-15 years of sharing computational models in life science. Nucleic Acids Res. 2020;48(D1):D407–D15. doi:10.1093/nar/gkz1055.

## References

1. Mardinoglu A, Agren R, Kampf C, Asplund A, Uhlen M, Nielsen J. Genome-scale metabolic modelling of hepatocytes reveals serine deficiency in patients with non-alcoholic fatty liver disease. Nature communications. 2014;5.

2. Ma H, Sorokin A, Mazein A, Selkov A, Selkov E, Demin O, et al. The Edinburgh human metabolic network reconstruction and its functional analysis. Mol Syst Biol. 2007;3:135. doi:10.1038/msb4100177. PubMed PMID: 17882155; PubMed Central PMCID: PMCPMC2013923.

3. Thiele I, Swainston N, Fleming RM, Hoppe A, Sahoo S, Aurich MK, et al. A community-driven global reconstruction of human metabolism. Nat Biotechnol. 2013;31(5):419–25. doi:10.1038/nbt.2488. PubMed PMID: 23455439; PubMed Central PMCID: PMCPMC3856361.

4. Trupp M, Altman T, Fulcher CA, Caspi R, Krummenacker M, Paley S, et al. Beyond the genome (BTG) is a (PGDB) pathway genome database: HumanCyc. Genome biology. 2010;11(1):O12.

5. Kanehisa M, Goto S, Sato Y, Kawashima M, Furumichi M, Tanabe M. Data, information, knowledge and principle: back to metabolism in KEGG. Nucleic acids research. 2013;42(D1):D199–D205.

6. Mardinoglu A, Agren R, Kampf C, Asplund A, Nookaew I, Jacobson P, et al. Integration of clinical data with a genome-scale metabolic model of the human adipocyte. Mol Syst Biol. 2013;9:649. doi:10.1038/msb.2013.5. PubMed PMID: 23511207; PubMed Central PMCID: PMCPMC3619940.

7. Cakir T, Patil KR, Onsan Z, Ulgen KO, Kirdar B, Nielsen J. Integration of metabolome data with metabolic networks reveals reporter reactions. Mol Syst Biol. 2006;2:50. doi:10.1038/msb4100085. PubMed PMID: 17016516; PubMed Central PMCID: PMCPMC1682015.

8. Patil KR, Nielsen J. Uncovering transcriptional regulation of metabolism by using metabolic network topology. Proceedings of the National Academy of Sciences of the United States of America. 2005;102(8):2685–9. doi:10.1073/pnas.0406811102.

